# Enhancing Pharmacometric Modeling with Full Bayesian Inference and Student’s t-Based M3 Censoring: A Simulated Population PK Study on Robust Handling of Outliers and Censored Data

**DOI:** 10.1101/2025.07.17.665409

**Authors:** Yiming Cheng, Yan Li

## Abstract

Pharmacometric modeling and simulation, with population pharmacokinetic (PopPK) modeling as a key component, play a crucial role in characterizing drug disposition and variability, supporting dose optimization and regulatory decision-making. Traditional maximum likelihood estimation (MLE) methods, although efficient, are sensitive to outliers, particularly when datasets contain observations below the limit of quantification (BLQ). As clinical datasets become increasingly complex, more robust modeling approaches are needed. In this study, a simulation-based evaluation was conducted to compare four modeling strategies combining different residual error structures (normal vs. Student’s t) and censoring methods (M1 vs. M3). Two-compartment pharmacokinetic profiles were simulated for fifty subjects, incorporating varying degrees of outlier contamination and BLQ data. Full Bayesian inference using Markov Chain Monte Carlo (MCMC) methods was employed to estimate posterior distributions of PK parameters at both the population and individual levels. Notably, the combination of Student’s t-distributed residuals with M3 censoring (Student’s t_M3) consistently produced the most accurate and precise parameter estimates across all simulation scenarios, even under extreme outlier contamination and substantial BLQ presence. The combination of Student’s t-distributed residuals with M3 censoring within a Bayesian framework offers a robust and resilient strategy for PopPK modeling, effectively addressing both outlier contamination and data censoring challenges. These findings support the broader adoption of robust Bayesian modeling techniques in pharmacometric practice, particularly for complex and irregular clinical datasets such as cell and gene therapies.

## Introduction

Pharmacometric modeling and simulation, with population pharmacokinetic (PopPK) modeling as a key component, provide critical tools for characterizing drug disposition and variability across patient populations. ^1,2^ These tools play an essential role in dose optimization and regulatory decision-making, especially in light of initiatives such as Project Optimus, launched by the FDA to improve dose selection strategies in oncology drug development. ^3–9^ Traditionally, maximum likelihood estimation (MLE) methods, implemented through gradient-based algorithms such as FO and FOCE in NONMEM®, have served as the standard approach and have been routinely used in the pharmacometric field for parameter estimation in PopPK and other PK/PD models. ^10,11^ These methods are computationally efficient and well-supported by asymptotic statistical theory, particularly when applied to large datasets with normally distributed residuals. However, two key issues frequently encountered in PopPK analysis are the presence of outliers and the handling of data below the lower limit of quantification (BLQ).

Outliers—observations markedly different from the general data trend—can distort parameter estimation, introduce bias, and compromise model stability. BLQ data, commonly observed in pharmacokinetic profiles, contain valuable information about drug elimination kinetics yet are often mishandled when treated as missing or excluded during modeling. ^12–18^ Current standard practices to address these challenges have limitations. Outlier handling typically involves subjective residual-based criteria, such as setting conditional weighted residuals (CWRES) thresholds for data exclusion. ^19^ Similarly, censored data are often managed through simple omission (M1 method), which can introduce bias into parameter estimates. Although the M3 method has been shown to offer a likelihood-based solution for BLQ data, ^12,13,17^ and robust error models like the Student’s t-distribution provide resilience to outliers, ^20–22^ their combined application within a unified modeling framework has not been systematically evaluated.

Recent advances in Bayesian computational methods, along with enhanced functionalities in pharmacometric software (e.g., NONMEM®’s GAMLN and various built-in probability density functions), ^10,11^ have now made it easy and practical to incorporate more sophisticated error structures and censoring techniques into PopPK analyses. In addition, Bayesian approaches provide full posterior distributions that capture both parameter estimates and their associated uncertainty without relying on asymptotic approximations. ^23,24^

The increasing complexity of clinical datasets, particularly in fields such as cell and gene therapies—where data variability is higher, increasing the chance of outlier observations, and assay methods such as qPCR for CAR-T therapies introduce a greater proportion of BLQ observations—further highlights the need for more robust modeling approaches to handle these combined challenges. ^25,26^ In such settings, combining Student’s t-distributed residuals with M3 censoring methods within a Bayesian framework may offer significant advantages over traditional approaches, enhancing the robustness and reliability of PopPK analyses even in the presence of substantial data irregularities.

The present study was designed to systematically investigate this approach. Using simulation scenarios that varied in both the severity of outlier presence (moderate and extreme) and the extent of data censoring, we evaluated the performance of four modeling strategies combining different residual error structures (normal vs. Student’s t) and censoring methods (M1 vs. M3). The goal of this work is to demonstrate that Bayesian modeling incorporating the Student’s t-based M3 censoring method can provide a robust and reliable framework for PopPK parameter estimation in challenging datasets, thereby advancing current pharmacometric modeling practices.

## Methods

### Data Simulation

To assess the effectiveness of full Bayesian inference combined with Student’s t-based M3 censoring for robust handling of outliers and censored data, a dedicated simulation study was conducted. Simulated datasets were generated and then fitted with various models to compare their performance in managing outliers and below-quantification-limit observations.

Specifically, intravenous (IV) two-compartment pharmacokinetic (PK) profiles were simulated for 50 virtual subjects using the following parameter values: clearance (CL) = 2 L/h; central compartment volume (V1) = 30 L; intercompartmental clearance (Q) = 5 L/h; and peripheral compartment volume (V2) = 80 L. Assuming a log-normal distribution for inter-individual variability (IIV) in PK parameters, an ETA of 20% was applied to all structural parameters, and a proportional residual error (σ) of 20% was assumed. No additional random effects or covariate effects were included. Each subject received a single 400 mg IV bolus dose, producing two distinct terminal phases, with a peak concentration (Cmax) of approximately 15 ng/mL and a concentration of about 2 ng/mL at approximately 15 hours post-dose, marking the α-to-β phase transition. Notably, an IV bolus was chosen purely for simplicity; the Student’s t_M3 likelihood is independent of the absorption model and can be applied unchanged to oral or other extravascular dosing schemes.

The PopPK dataset used for model fitting was generated by selecting specific timepoints from the simulated data. The sampling timepoints (0, 4, 8, 12, 16, 20, 30, 40, 50 and 60 hours) were selected to adequately characterize the two-compartment PK model based on the simulated data, allowing reliable and precise parameter estimation in the absence of outliers and BLQ data during model fitting. These reference estimates were then used to assess method performance under various outlier and BLQ scenarios, as outlined below.

Based on these profiles, four lower limits of quantification (LLOQ = 0.90, 1.13, 1.37, and 1.60 ng/mL) were applied to create datasets, representing the range of BLQ rates from 10% to 40%, which reflects increasing degrees of censoring. For each LLOQ, two censoring approaches were prepared: M1, in which values below the LLOQ are treated as missing; and M3, in which values below the LLOQ are incorporated into the likelihood as censored data. To introduce outliers, one or two concentration measurements in the PK profiles of a subset of subjects (representing 10%, 45%, or 80% of the cohort) were selected from either the early α-terminal phase or the later β terminal phase and multiplied by factors of 1 (baseline), 5.5 (450% increase), or 10 (900% increase), thereby generating no, moderate, and extreme deviations, respectively. This approach preserves the underlying PK trajectory while imposing controlled, phase-specific outliers for robust method evaluation.

Each dataset—defined by its LLOQ threshold and outlier severity—was fitted under four model configurations: normal residual error with M1 censoring (Model #1); normal residual error with M3 censoring (Model #2); Student’s t residual error with M1 censoring (Model #3); and Student’s t residual error with Student’s t-based M3 censoring (Model #4).

Of note, no clinical data were analyzed in this study; all results are based on *in silico* datasets generated exclusively for methodological comparison.

## PopPK Model with Full Bayesian Approach

In the current study, population pharmacokinetic (PopPK) analysis was conducted using the nonlinear mixed-effects modeling software NONMEM® (Version 7.5.0, ICON Development Solutions, North Wales, PA, USA), applying a full Bayesian framework through Markov Chain Monte Carlo (MCMC) methods. ^10,11^ Data preparation, model initialization, execution, and post-processing were all performed in R.

In this full Bayesian MCMC approach, pharmacokinetic parameters were modeled as random variables with prior probability distributions, in contrast to traditional methods where parameters are treated as fixed unknowns. Rather than seeking point estimates for θ (fixed effects), σ (residual variability), and Ω (inter-individual variability) using traditional gradient descent-based algorithms such as FO or FOCE, MCMC Bayesian analysis aims to generate large samples of parameter sets from the posterior distribution. These posterior samples support statistical inference by enabling the calculation of summary measures such as means, standard deviations, and credible intervals for population parameters.

In addition, unlike maximum likelihood–based analyses, the Bayesian method typically does not compute an objective function value. Instead, MCMC sampling constructs the joint posterior distribution over the entire parameter space. The MCMC process was divided into two distinct phases: an initial burn-in phase, characterized by directional trends in likelihood and parameter estimates, and thus unsuitable for statistical summarization, and a stationary sampling phase, where parameter values fluctuate randomly without systematic drift. In the present study, the number of iterations was set to 1,500 for the burn-in phase and 4,000 for the stationary phase, with only samples from the stationary phase used for posterior inference.

Specification of prior information was a critical element of the Bayesian model. In the current study, uninformative priors were assigned throughout: multivariate normal distributions with large variances were used for fixed effects, and inverse Wishart distributions were applied for covariance structures governing random effects. Residual variability was modeled either by a normal distribution or, in some cases, by a Student’s t-distribution, depending on the model configuration.

Posterior samples generated from the stationary phase were used to derive individual- and population-predicted concentrations, as well as to characterize posterior parameter distributions. A log-link model was applied throughout to stabilize residual variability across the concentration range. These outputs were employed for both model diagnostic evaluation and statistical inference regarding the underlying pharmacokinetic behavior.

## PopPK Modeling with Normal or Student’s t-Based M1/M3 Censoring Methods

The simulated datasets were analyzed using the same two-compartment PK model structure that was used for their generation. This model included parameters for clearance (CL), central compartment volume (V1), intercompartmental clearance (Q), and peripheral compartment volume (V2), with inter-individual variability (IIV) assumed to follow a log-normal distribution for all PK parameters.

The observed data were modeled directly through the -2 × log-likelihood of the data, using either a normal distribution or a Student’s t distribution to characterize the residual error structure. Both observed and model-predicted concentrations were log-transformed prior to model fitting to ensure consistent error scaling across the concentration range. Specifically, the residual error model was defined by a normal distribution (Equation 1) or a Student’s t distribution (Equation 2):

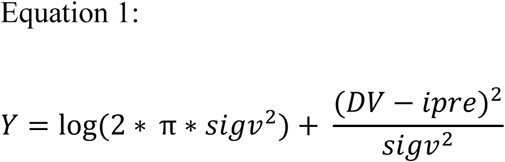

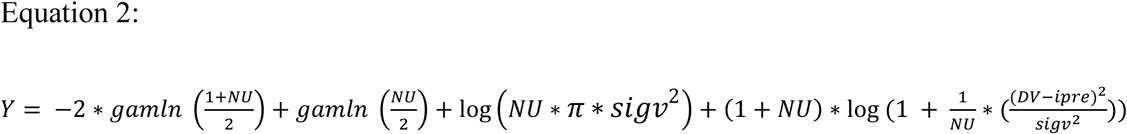

where Y refers to -2 × log-likelihood of the data, gamln refers to the NONMEM® built-in function that accurately calculates the logarithm of the gamma function, NU denotes the degrees of freedom for the Student’s t distribution, σ (sigv) represents the standard deviation of the residual distribution, DV corresponds to the log-transformed observed concentration, and IPRE corresponds to the log-transformed model-predicted concentration.

For censored observations (i.e., concentrations below the lower limit of quantification), different methods were applied depending on the residual error model. For models using a normal residual distribution (Model #2), the NONMEM® built-in function PHI() was used to compute the cumulative probability for censored data based on the standard normal distribution. For models using a Student’s t residual distribution (Model #3), the STUDENTTCDF() function was used to appropriately account for censoring under the Student’s t framework.

Model evaluation focused on both parameter recovery and predictive performance. Estimated PK parameters from each model were compared against the true values originally used for simulation to assess bias and recovery accuracy. Posterior predictive distributions were generated for each model, and the corresponding 90% credible intervals were evaluated to assess bias and precision across models. In addition, model-predicted concentration–time profiles were constructed and compared against the original simulation-based profiles, providing a visual and quantitative assessment of model performance. Comparative evaluations were performed across all four model configurations (Models #1, #2, #3, and #4) to identify the best-performing method under scenarios involving outliers and censored data.

## Results

### Simulated PK Profiles

Figure 1 presents the simulated PK profiles for 50 virtual participants. A typical two-phase decline was observed, with a reflection time point at approximately 15 hours post-dose and a corresponding concentration of around 2 ng/mL. The earlier α phase reflected the rapid distribution phase, while the later β phase corresponded to the slower elimination phase.

**Figure 1.**
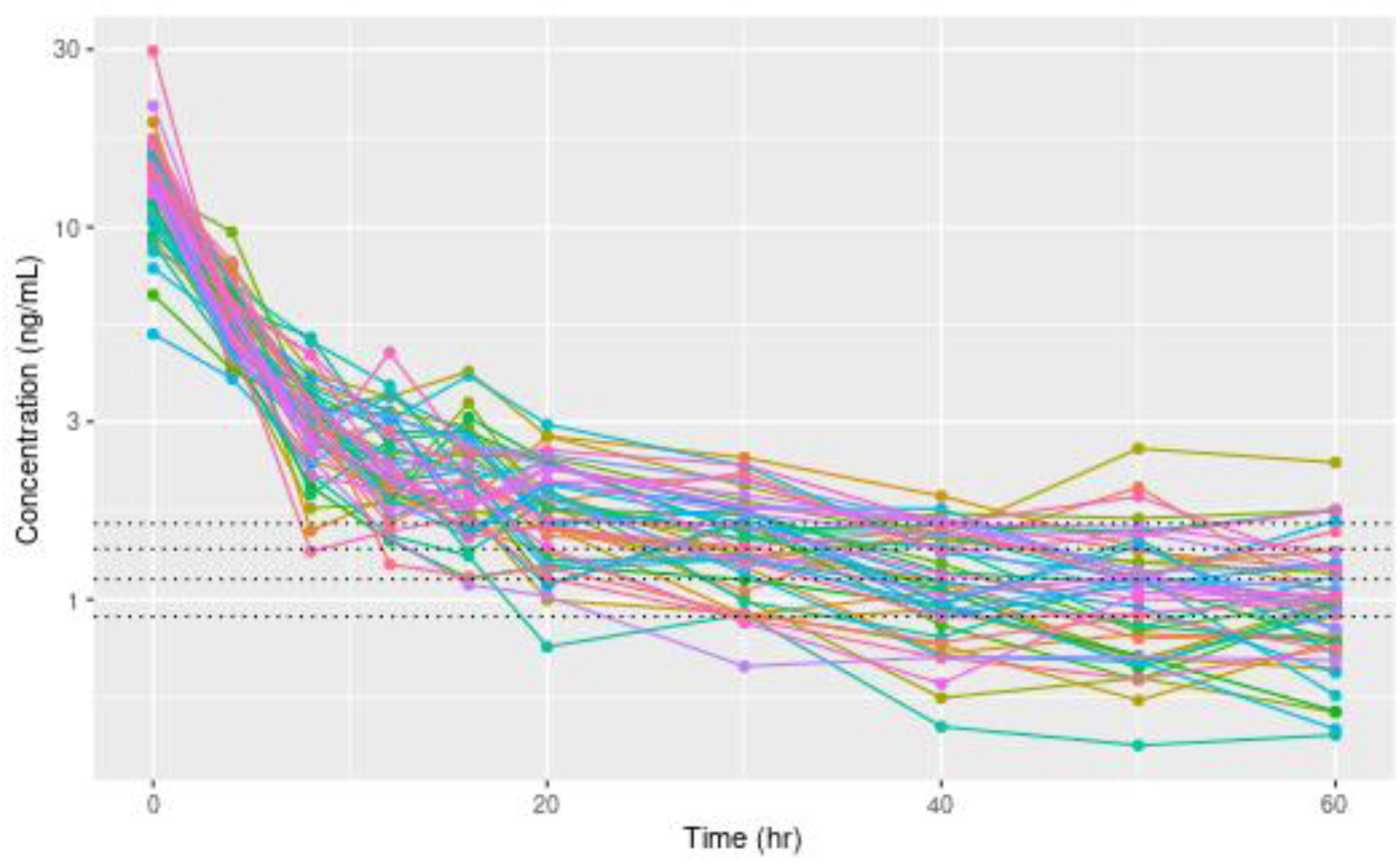
Simulated concentration–time profiles.

Four dotted black lines indicate the applied lower limits of quantification (LLOQ = 0.90, 1.13, 1.37, and 1.60 ng/mL), representing increasing degrees of censoring across the concentration range. Of note, the LLOQ values were intentionally selected near the intersection of the α and β phases, where the transition between the two phases occurs. This design allowed for a focused evaluation of how well different modeling methods handle censored data near critical pharmacokinetic transitions. In particular, the analysis assessed the impact of different LLOQ thresholds on the estimation of parameters governing the β-terminal phase, such as clearance, and volume of peripheral compartment, which are sensitive to late-phase concentration data.

## Impact of LLOQ Cutoffs on PK Parameter Estimation Across Modeling Methods

Table 1 presents the true values and population pharmacokinetic parameter estimates derived from the posterior distributions for all four methods, based on a specific scenario where the multiplicative factor is 10 and the LLOQ cutoff is 1.6 ng/mL.

**Table 1.**
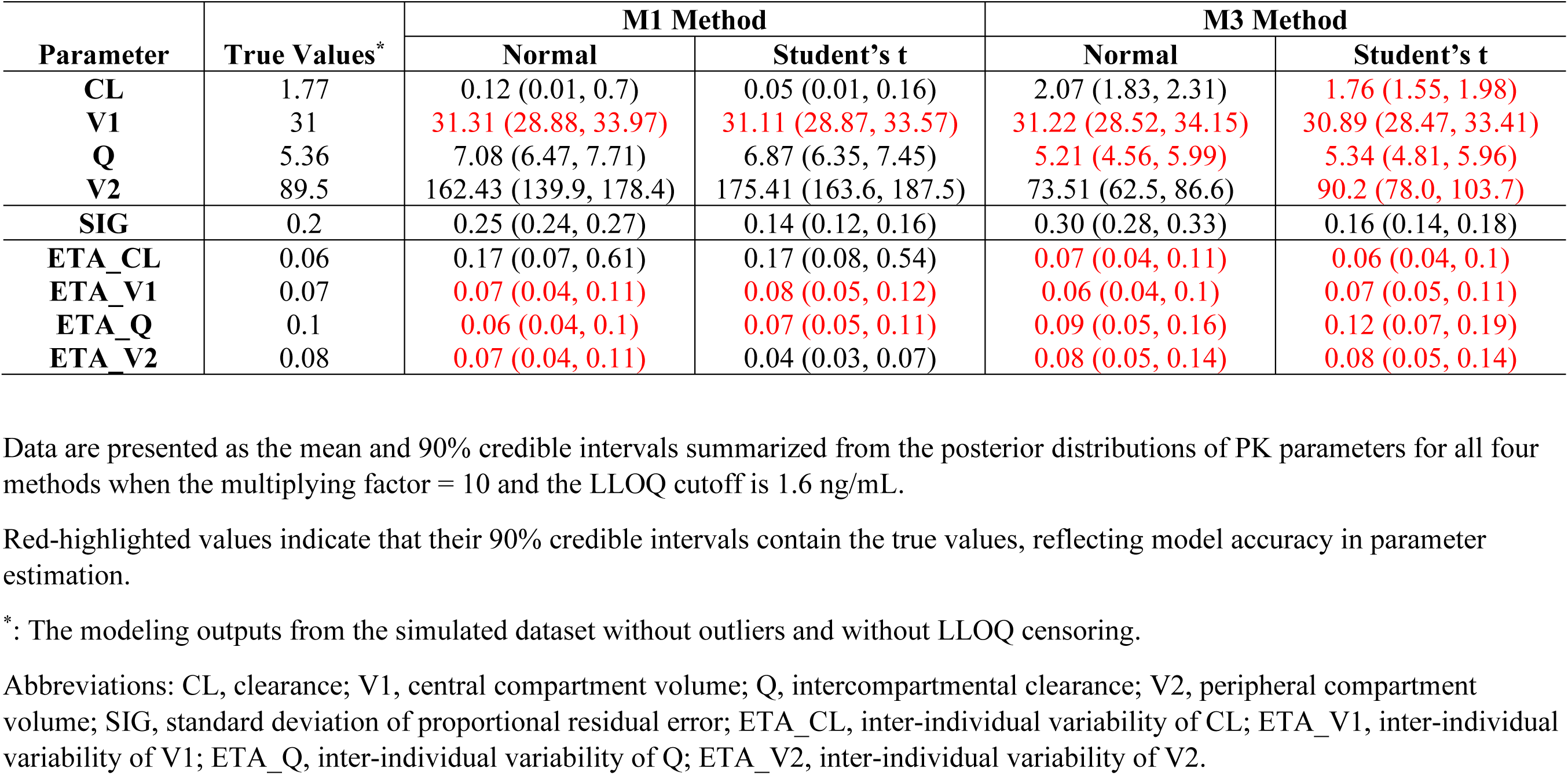
True PopPK parameter values and parameter estimates summarized from the posterior distributions.

| Parameter | True Values* | M1 Method |  | M3 Method |  |
| --- | --- | --- | --- | --- | --- |
|  |  | Normal | Student's t | Normal | Student's t |
| CL | 1.77 | 0.12 (0.01, 0.7) | 0.05 (0.01, 0.16) | 2.07 (1.83, 2.31) | 1.76 (1.55, 1.98) |
| V1 | 31 | 31.31 (28.88, 33.97) | 31.11 (28.87, 33.57) | 31.22 (28.52, 34.15) | 30.89 (28.47, 33.41) |
| Q | 5.36 | 7.08 (6.47, 7.71) | 6.87 (6.35, 7.45) | 5.21 (4.56, 5.99) | 5.34 (4.81, 5.96) |
| V2 | 89.5 | 162.43 (139.9, 178.4) | 175.41 (163.6, 187.5) | 73.51 (62.5, 86.6) | 90.2 (78.0, 103.7) |
| SIG | 0.2 | 0.25 (0.24, 0.27) | 0.14 (0.12, 0.16) | 0.30 (0.28, 0.33) | 0.16 (0.14, 0.18) |
| ETA_CL | 0.06 | 0.17 (0.07, 0.61) | 0.17 (0.08, 0.54) | 0.07 (0.04, 0.11) | 0.06 (0.04, 0.1) |
| ETA_V1 | 0.07 | 0.07 (0.04, 0.11) | 0.08 (0.05, 0.12) | 0.06 (0.04, 0.1) | 0.07 (0.05, 0.11) |
| ETA_Q | 0.1 | 0.06 (0.04, 0.1) | 0.07 (0.05, 0.11) | 0.09 (0.05, 0.16) | 0.12 (0.07, 0.19) |
| ETA_V2 | 0.08 | 0.07 (0.04, 0.11) | 0.04 (0.03, 0.07) | 0.08 (0.05, 0.14) | 0.08 (0.05, 0.14) |

Data are presented as the mean and 90% credible intervals summarized from the posterior distributions of PK parameters for all four methods when the multiplying factor = 10 and the LLOQ cutoff is 1.6 ng/mL.
Red-highlighted values indicate that their 90% credible intervals contain the true values, reflecting model accuracy in parameter estimation.
Abbreviations: CL, clearance; V1, central compartment volume; Q, intercompartmental clearance; V2, peripheral compartment volume; SIG, standard deviation of proportional residual error; ETA\_CL, inter-individual variability of CL; ETA\_V1, inter-individual variability of V1; ETA\_Q, inter-individual variability of Q; ETA\_V2, inter-individual variability of V2.

Figure 2 presents the posterior distributions of the population PK parameter estimates from the full Bayesian approach generated by the four models when one outlier was created from a concentration point within the β-terminal phase. Column panels represent datasets generated using different LLOQ cutoffs (LLOQ = 0.90, 1.13, 1.37, and 1.60), while row panels represent different multiplicative factors used for outlier creation (factors = 1, 5.5, and 10); larger factors reflect more extreme deviations. The dotted black lines indicate the true parameter values used in the original simulation.

**Figure 2.**
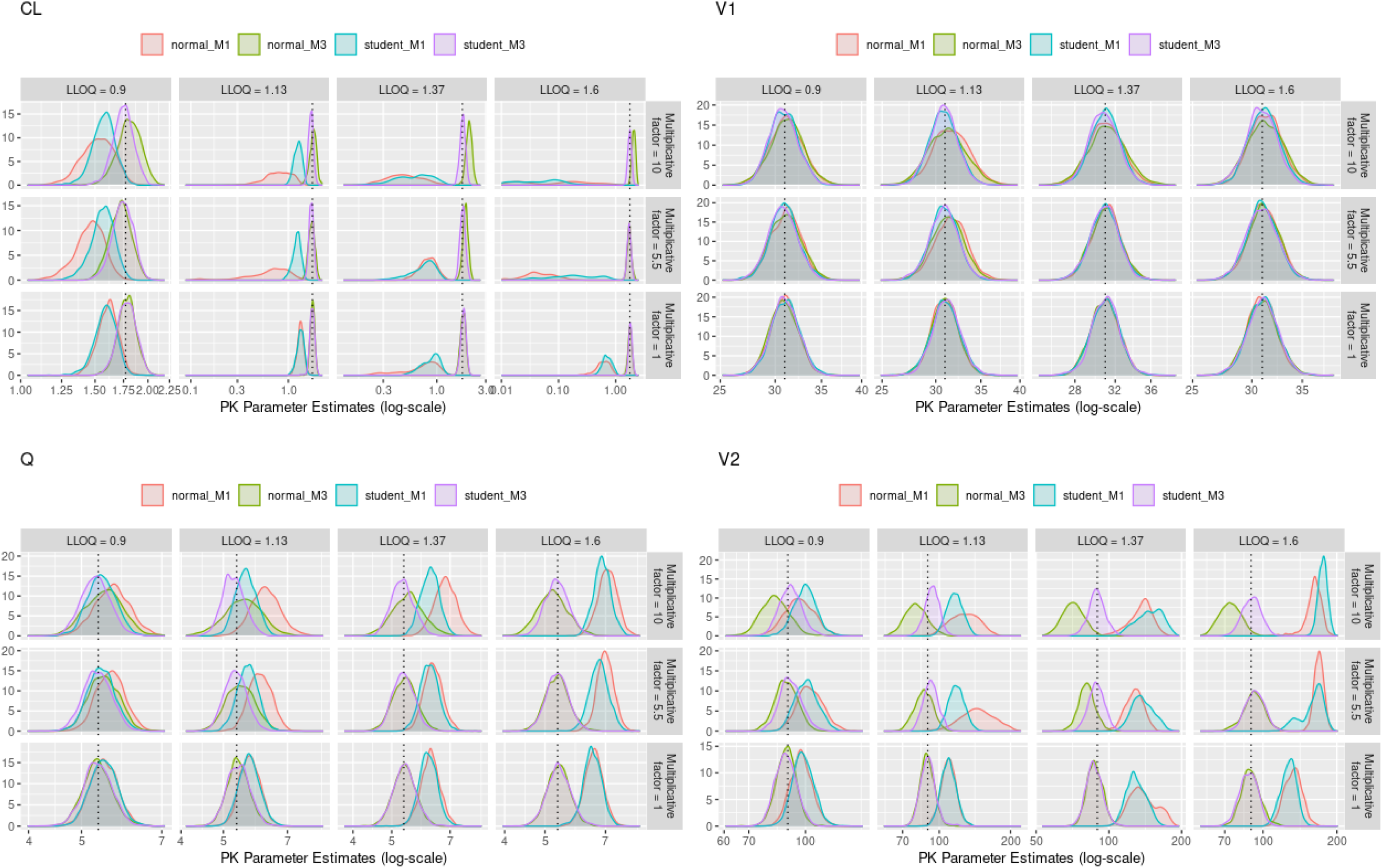
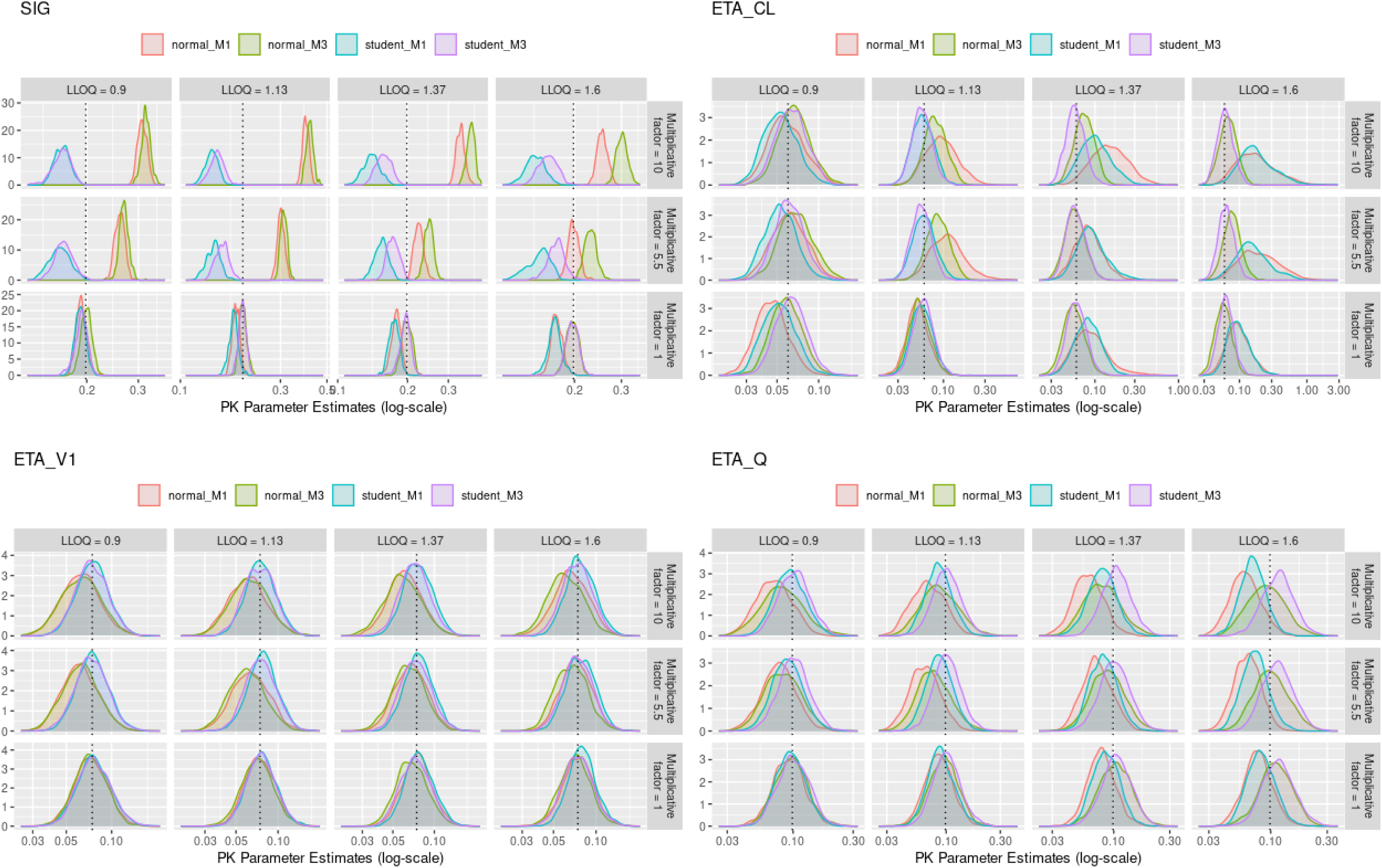
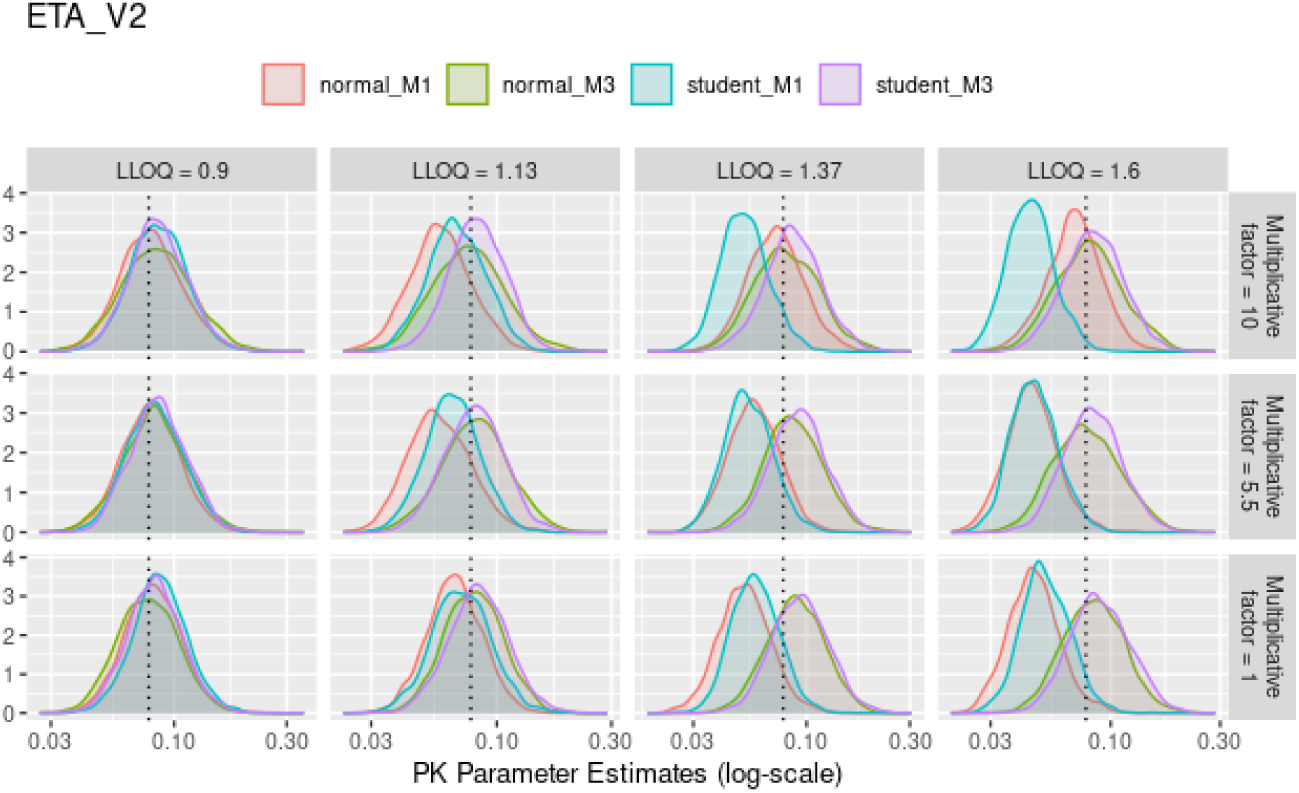
Posterior distributions of population PK parameter estimates from the full Bayesian approach. Dotted black lines indicate the true values (i.e., the modeling outputs from the simulated dataset without outliers and without LLOQ censoring). Column panels show the posterior distributions of PK parameter estimates under different LLOQ cutoffs (LLOQ = 0.9, 1.13, 1.37, and 1.6 ng/mL). Row panels show the posterior distributions under different multiplicative factors (factors = 1, 5.5, and 10) used for outlier generation. CL: clearance; V1: central compartment volume; Q: intercompartmental clearance; V2: peripheral compartment volume; SIG: standard deviation of proportional residual error; ETA_CL: inter-individual variability of CL; ETA_V1: inter-individual variability of V1; ETA_Q: inter-individual variability of Q; ETA_V2: inter-individual variability of V2.

In the bottom panels, corresponding to a multiplying factor of 1 (i.e., no outlier introduced), it is evident that as the LLOQ cutoff increases, the M3 methods—whether based on the normal or Student’s t distribution—maintain accurate estimations of CL, Q, and V2, with good precision as demonstrated by narrow posterior distributions. For V1, all four modeling methods produced similarly accurate estimates with comparable precision, which is expected given that V1 is primarily informed by early time points in the concentration–time profile and is therefore less affected by data censoring or outliers located in the β-terminal phase.

Additionally, due to the treatment of censored data as missing, both the normal_M1 and Student’s t_M1 methods showed a tendency to underestimate residual variability. This, in turn, introduced biases in the estimation of inter-individual variability as the LLOQ cutoff increased, manifesting as overestimation for CL and underestimation for Q and V2, while having no notable impact on the inter-individual variability of V1.

In summary, in the absence of outliers, both the normal_M3 and Student’s t_M3 methods provided accurate and precise PK parameter estimates, demonstrating robustness across varying LLOQ thresholds. These findings are consistent with prior conclusions regarding the robustness and efficiency of M3 methods in handling censored LLOQ data.

## Impact of Outliers on PK Parameter Estimation Across Modeling Methods

The middle and top panels of Figure 2 correspond to scenarios where multiplying factors of 5.5 and 10 were applied, representing moderate and extreme outlier deviations, respectively. For clearance (CL), both the normal_M3 and Student’s t_M3 methods continued to provide accurate and precise parameter estimates, with posterior distributions closely centered around the true values and exhibiting relatively narrow density profiles. However, under extreme outlier conditions, the normal_M3 method showed overestimation of CL compared to the Student’s t_M3 method, indicating reduced robustness of the normal_M3 approach in handling severe outlier situations.

At moderate outlier levels (multiplying factor of 5.5), the Student’s t_M1 method also produced reasonably precise CL estimates, although slight bias was observed relative to the true value. In contrast, the normal_M1 method failed to produce an accurate CL estimate, demonstrating clear sensitivity to the influence of moderate outliers.

For the intercompartmental clearance (Q), all four modeling methods produced estimates with comparable precision, as indicated by similar posterior distribution widths. However, both the normal_M3 and Student’s t_M3 methods again showed better accuracy, providing estimates closer to the true values compared with the normal_M1 and Student’s t_M1 methods.

Notably, for the volume of distribution of the peripheral compartment (V2)—the parameter most sensitive to the characterization of the β-terminal phase—the Student’s t_M3 method consistently provided the most accurate and precise parameter estimates across all conditions.

Even under extreme conditions—when an outlier was created using a multiplying factor of 10 combined with a high LLOQ cutoff of 1.60 ng/mL, resulting in substantial data censoring—the Student’s t_M3 method maintained strong performance, outperforming all other modeling strategies. This was demonstrated by the fact that, for all PK parameters (CL, V1, Q, and V2), the 90% credible intervals derived from the posterior distributions contain the true values. In contrast, although the normal_M3 method produced more accurate parameter estimates than the normal_M1 and Student’s t_M1 methods, its 90% credible intervals did not consistently contain the true values, as shown in Table 1.

In summary, in the presence of moderate to extreme outliers, the Student’s t_M3 method demonstrated superior resilience, delivering accurate and precise PK parameter estimates across a wide range of outlier severities and censoring conditions. These results highlight the robustness of the Student’s t_M3 approach for simultaneously handling both outliers and censored data in PK modeling.

## Impact of Outlier Position and Outlier Prevalence on PK Parameter Estimation Across Modeling Methods

Additional simulation and modeling analyses were conducted to further evaluate the robustness of different modeling methods. The scenarios examined included: (1) one outlier created from a concentration point within the early α-terminal phase; (2) two outliers created, one from the early α-terminal phase and one from the late β-terminal phase; (3) two outliers created from two concentration points within the early α-terminal phase; and (4) two outliers created from two concentration points within the late β-terminal phase. Across all of these scenarios, the Student’s t_M3 method consistently outperformed the other three methods in handling both outliers and censored data simultaneously (Supplementary Figures S4-S7).

In addition, the impact of varying the proportion of subjects with outliers on PK parameter estimation was evaluated. It was observed that even when only 10% of subjects (5 out of 50) had outliers, there was a significant effect on both population-level and individual-level PK parameter estimates. Among all methods, only the Student’s t distribution combined with the M3 approach demonstrated sufficient robustness to maintain accurate and precise estimation in the presence of outlier contamination (Supplementary Figures S4-S7).

## Impact of LLOQ Cutoffs and Outliers on Individual PK Parameter Estimation Across Modeling Methods

Figures 3 and 4 present the posterior distributions of PK parameter estimates and corresponding concentration–time profiles for two representative subjects. Consistent with the findings from the population PK parameter analysis, for Subject ID = 17, the presence of outliers at 16 and 20 hours caused distortion in the profiles generated from the normal distribution–based parameter estimates (regardless of M1 or M3 methods, shown by the red and green curves). These profiles bent upward to accommodate the influence of the outliers on the likelihood. In contrast, the profiles generated from the Student’s t-based parameter estimates (blue and purple curves for the M1 and M3 methods, respectively) remained well aligned with the simulated true profiles, reflecting the resilience of the Student’s t distribution to outlier effects.

**Figure 3.**
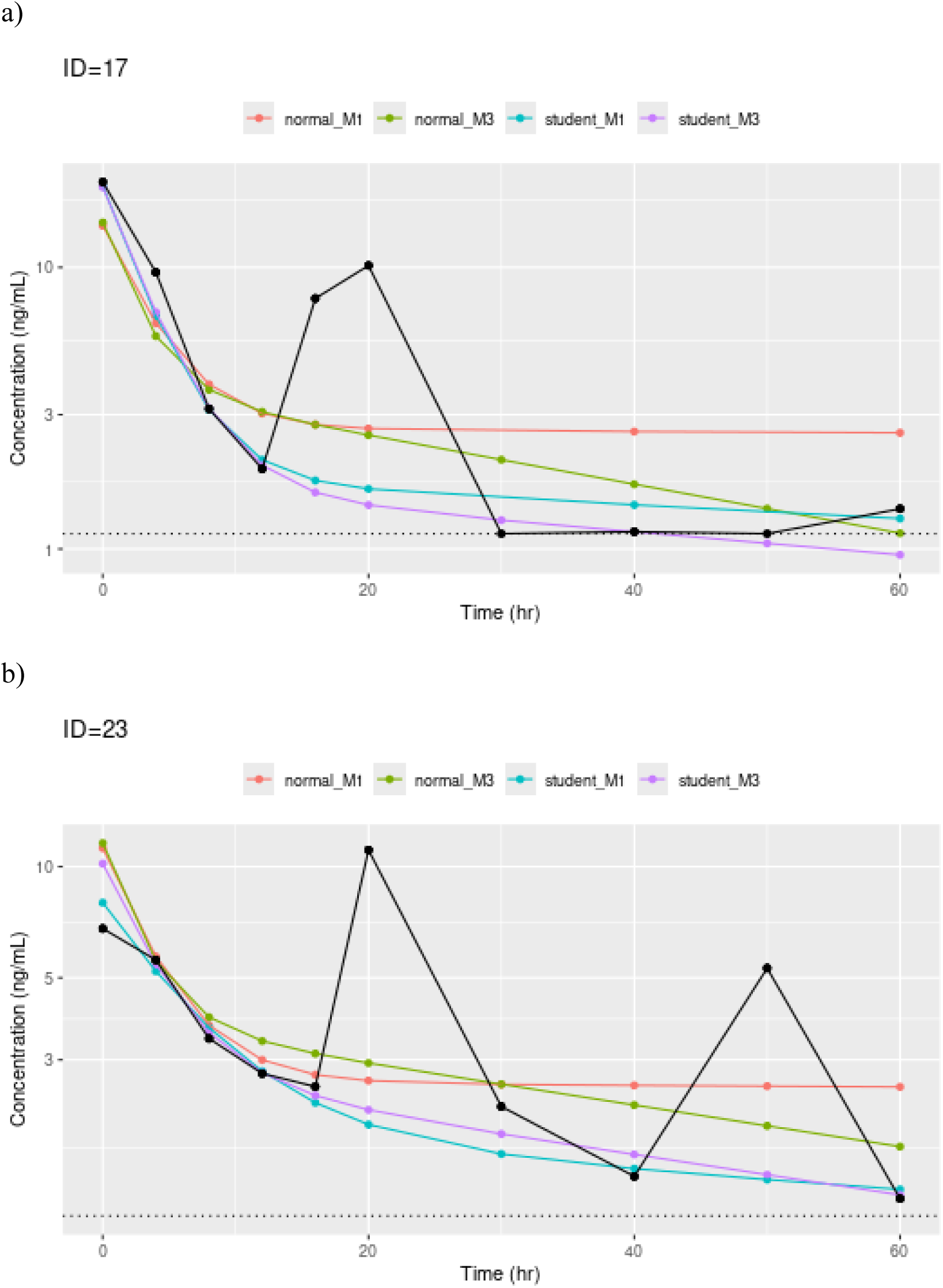
Concentration–time profiles from the full Bayesian approach for two selected subjects. Black lines and symbols represent the observed data. Colored lines and symbols represent profiles generated from the fitted parameters obtained using different modeling methods.

**Figure 4.**
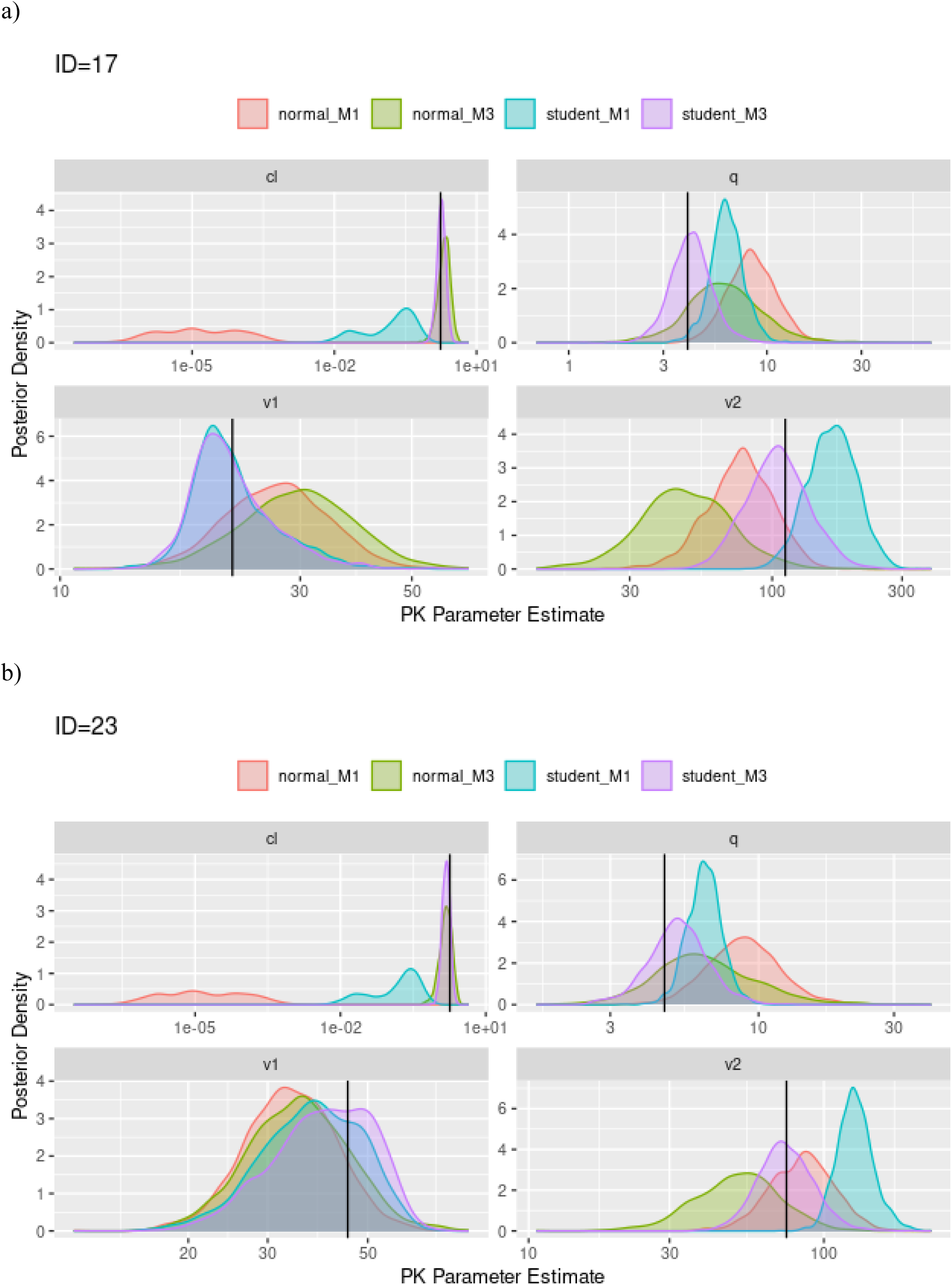
Posterior distributions of individual PK parameter estimates from the full Bayesian approach for two selected subjects. Black lines indicate the true values used in simulations.

As a result, the posterior density distributions of all four PK parameters estimated by the Student’s t_M3 method were not only accurate but also relatively precise, as demonstrated by narrower posterior density profiles compared to the other methods. For the normal_M3 method, the incorporation of censored data improved the capture of the β-terminal phase compared with the normal_M1 method (green curve versus red curve). Similarly, by incorporating censored data, the Student’s t_M3 method outperformed the Student’s t_M1 method (purple curve versus blue curve). Similar findings were also observed for Subject ID = 23, further reinforcing these findings.

Beyond these two subjects, the accuracy of individual PK parameter estimates for all subjects across the four methods in the analysis underlying Table 1 was calculated and presented in Supplementary Figure S3. The results showed that the Student’s t_M3 method consistently demonstrated the highest accuracy across all individuals, supporting its ability to provide reliable parameter estimates at both the population and individual levels.

In summary, consistent with the population-level results, combining both the Student’s t distribution and the M3 method led to more accurate and precise individual PK parameter estimates across most subjects.

## Discussions and Conclusions

Traditional gradient descent–based Maximum Likelihood Estimation (MLE) methods remain a cornerstone of pharmacometric modeling, particularly in population pharmacokinetic (PopPK) and PK/PD analyses. ^1,2,27^ Their computational efficiency and well-established theoretical foundations have long made them the preferred approach across clinical pharmacology and pharmacometric communities. Implementations such as the FO and FOCE methods in NONMEM® and other commercial software are widely regarded as industry gold standards. ^10,11^ However, as typical frequentist approaches, MLE methods inherently focus on generating point estimates for population parameters and generally require additional procedures—such as Fisher Information Matrix–based standard error calculations, bootstrapping, or likelihood profiling—to assess parameter uncertainty and to support further statistical inference. ^1,2^

In contrast, simulation/sampling-based full Bayesian inference provides a much richer output. ^24^ Rather than relying solely on point estimates, Bayesian methods generate complete posterior distributions for model parameters through Gibbs sampling or Metropolis-Hastings algorithms. ^10,11^ This allows direct and simultaneous characterization of both parameter estimates and their associated uncertainty, making subsequent statistical inference more straightforward. In the current simulation study, a full Bayesian framework was utilized to illustrate these advantages. Through Markov Chain Monte Carlo (MCMC) sampling, posterior distributions were obtained for each PK parameter (CL, V1, Q, and V2) at both the population and individual subject levels under various scenarios of data censoring and outlier presence. These posterior distributions enabled direct derivation of credible intervals, providing intuitive and comprehensive statistical summaries without the need for additional asymptotic approximations. We further compared full Bayesian inference (via MCMC) with traditional MLE using NONMEM’s FOCEI method across the same four model configurations. While point estimates were generally consistent between approaches (Supplementary Table S1), the MLE method frequently failed in most configurations. As a result, standard errors and confidence intervals could not be obtained without additional resampling procedures (e.g., bootstrap). These issues highlight the known numerical sensitivity of gradient-based MLE methods when applied to datasets with both severe outliers and extensive BLQ censoring. In this study, the posterior density profiles for parameter estimates at both the population and individual levels offered a straightforward assessment of parameter uncertainty, with narrower and more sharply peaked distributions indicating higher precision in the parameter estimates. Although full Bayesian analysis has historically seen limited application in the PK/PD field, partly due to intensive computational demands, advances in hardware and software platforms have now made Bayesian approaches increasingly practical. ^10,11^ Our analysis, performed using parallel computing strategies in R and NONMEM®, demonstrated that even complex Bayesian models can be efficiently fitted within a matter of minutes.

Outliers—data points that deviate markedly from expected trends—are an inevitable feature of real-world PK datasets, arising from sources such as assay variability, protocol deviations, or data entry errors. ^19^ Traditional normal-distribution–based residual error models are highly sensitive to such aberrant observations, often resulting in biased parameter estimates or inflated uncertainty. Our previous work demonstrated the advantages of utilizing the Student’s t-distribution to model residual variability in a real-world case study. ^21^ In the current study, even a small proportion of outliers (e.g., affecting only 10% of subjects) significantly impacted parameter recovery when using normal-based models, particularly under the M1 approach, which treats censored data simply as missing. To mitigate the influence of outliers, a common practice in the pharmacometric field is to apply a cutoff based on conditional weighted residuals (CWRES); however, this approach remains subjective. Setting a lower CWRES threshold may result in the exclusion of valuable data, whereas setting a higher threshold risks retaining influential outliers, potentially biasing the final PK parameter estimates.

The Student’s t-distribution offers a heavier-tailed alternative to the normal distribution, providing inherent resilience against the influence of extreme values. ^22^ This robustness arises because large residuals contribute less weight to the likelihood compared to the normal model. Our results clearly demonstrate this advantage: models incorporating Student’s t-distributed residual errors (both Student’s t_M1 and Student’s t_M3) consistently produced more stable and accurate PK parameter estimates across a range of outlier severities. Even under extreme conditions, Student’s t-based models maintained close alignment with the true simulated parameter values. In particular, the Student’s t methods exhibited remarkable stability, with narrow posterior distributions and corresponding credible intervals that consistently contain the true values, all without the need for subjectively setting a CWRES cutoff. These findings reinforce the prior literature supporting the use of the Student’s t-distribution for robust statistical modeling and highlight its underutilized potential within the pharmacometrics field.

Data below the lower limit of quantification (LLOQ)—a type of censored data—are another inevitable feature of real-world PK datasets. These are values that fall below the lowest concentration at which a bioanalytical method has been validated to produce results with acceptable accuracy and precision. It is well recognized that such data contain valuable information for PK parameter estimation. Therefore, rather than removing them from the analysis (as in the M1 method), various censoring strategies have been developed. ^12,13,15–18^ Among these, the normal-distribution–based M3 method, which properly integrates censored observations into the likelihood function as interval-censored data rather than discarding them as missing, has been shown to be the most powerful approach for improving parameter estimation. ^12,13,17^ Preserving information from data points falling below the LLOQ is particularly important in PK studies, where late-phase samples provide critical information about drug elimination kinetics.

In the present study, we report for the first time the combination of M3 censoring methods with Student’s t-distributed residuals, achieving the highest levels of robustness and accuracy observed to date. When combined with the outlier-handling strength of the Student’s t error model, the resulting Student’s t_M3 configuration consistently produced superior parameter estimation across all scenarios evaluated. Specifically, under moderate and extreme outlier conditions with high LLOQ cutoffs, the Student’s t_M3 method produced the most accurate and precise estimates for CL, V1, Q, and V2, as demonstrated by posterior distributions closely centered around the true simulation values and narrow credible intervals. Even when multiple outliers were present, or when substantial data censoring occurred (e.g., with LLOQ set near the α/β phase transition), the Student’s t_M3 method preserved model integrity and reliability.

In contrast, methods based on normal residuals, even when combined with M3 censoring (Model #2), showed marked degradation in performance under these challenging conditions. Overall, the combination of Student’s t-distributed residuals and M3 censoring effectively addresses two major challenges simultaneously—outlier contamination and LLOQ censoring—making it a highly powerful strategy for PopPK model development and inference.

Our simulation study offers findings highly relevant to the pharmacometric field, particularly given the increasing complexity of clinical datasets. Fields such as oncology, cell therapies, and biologics often exhibit greater data variability, increasing the likelihood of outlier observations, while assay methods such as qPCR for CAR-T therapies introduce a higher proportion of BLQ observations—further highlighting the need for more robust modeling approaches to handle these combined challenges. More specifically, CAR-T therapies have become increasingly popular in recent years. Because CAR-T assets are living cells—viable drugs—their PK profiles exhibit a higher frequency of outliers compared to traditional PK profiles from small molecules or protein-based therapeutics. ^25,26^ Furthermore, CAR-T pharmacokinetics are often characterized using a unique bioanalytical method, notably polymerase chain reaction (PCR) assay, which tend to produce a greater proportion of below the limit of quantification (BLQ) data during PK assessments. Given these characteristics, applying the Student’s t-based M3 method to characterize the PK properties of CAR-T therapies is highly relevant and currently under active investigation within our group. Our findings support a broader shift in PopPK modeling practices toward routinely adopting robust error models and likelihood-based handling of censored data, rather than relying solely on traditional normal residuals and subjective exclusion rules based on CWRES thresholds. With the increased accessibility of full Bayesian modeling frameworks and the capability to implement Student’s t-based error structures in modern software platforms (e.g., using the GAMLN and STUDENTTCDF functions in NONMEM®), there are now few technical barriers to adopting these more robust approaches in routine pharmacometric practice.

Of note, while the Student’s t-based M3 approach provided optimal performance in the current study—it is not without practical and theoretical caveats. First, although a heavy-tailed residual distribution offers strong protection against influential observations, it can reduce statistical efficiency when the data are approximately normal and true outliers are absent, resulting in slightly wider credible intervals than a well-specified normal model. In addition, precise estimation of the Student’s-t degrees-of-freedom (ν) parameter can also be challenging in sparse-sampling designs or when the proportion of BLQ data is high; in such situations, informative priors or sensitivity analyses are advisable. Finally, the MCMC implementation of the Student’s t_M3 model is more computationally intensive than its normal-based counterpart and requires convergence diagnostics that may be unfamiliar to some practitioners. These considerations should be weighed against the method’s robustness benefits when selecting an analysis strategy for a given dataset.

In conclusion, this study systematically evaluates full Bayesian PopPK modeling integrating Student’s t-distributed residual error models and M3-based censoring. Three key findings emerge: (1) full Bayesian inference yields richer and more informative outputs compared to traditional MLE methods; (2) Student’s t-distributed error models demonstrate superior resilience to outlier contamination relative to normal-based models; and (3) the combination of Student’s t residuals with M3 censoring delivers a powerful and robust framework for handling challenging PK datasets characterized by both outliers and censored observations. These findings support the broader adoption of Bayesian methods and robust error modeling strategies to enhance the accuracy, reliability, and scientific rigor of PopPK analyses in drug development.

Future work will focus on extending this framework to real-world datasets, including applications in characterizing pharmacokinetics of cell therapies.

## Consent for publication

All the authors have reviewed and concurred with the manuscript.

## Authors’ contributions

Y.C. and Y.L. contributed to conception and design; Y.C. and Y.L. contributed to acquisition of data; Y.C. and Y.L. contributed to analysis; all authors contributed to interpretation of data; Y.C. and Y.L. drafted and revised the article. Both authors made substantial contributions to conception and design, acquisition of data, or analysis and interpretation of data; took part in drafting the article or revising it critically for important intellectual content; agreed to submit to the current journal; gave final approval of the version to be published; and agreed to be accountable for all aspects of the work.

## Acknowledgments

The authors thank Robert J. Bauer from ICON for his assistance with the use of the NONMEM built-in density functions.

