## supplementary for "Enhancing Pharmacometric Modeling with Full Bayesian Inference and Student’s t-Based M3 Censoring: A Simulated Population PK Study on Robust Handling of Outliers and Censored Data"

**Supplementary Table S1.** True Population PK Parameter Values and Corresponding Estimates from MLE Methods.

| Parameter | True Values* | M1 Method | | M3 Method | |
|---|---|---|---|---|---|
| | | Normal | Student's t | Normal | Student's t |
| **CL** | 1.77 | 0.16 (0.13, 0.2) | 0.88 | 2.13 | 2.0 |
| **V1** | 31 | 31.68 (11.59, 86.56) | 30.58 | 31.06 | 30.23 |
| **Q** | 5.36 | 7.14 (6.77, 7.53) | 6.41 | 5.33 | 5.34 |
| **V2** | 89.5 | 149.1 (136.98, 162.28) | 118.3 | 72.54 | 78.75 |
| **SIG** | 0.2 | 0.28 (0.25, 0.31) | 0.16 | 0.31 | 0.18 |
| **ETA_CL** | 0.06 | 0.79 (0.79, 0.79) | 0.03 | 0.04 | 0.05 |
| **ETA_V1** | 0.07 | 0.02 (0.02, 0.02) | 0.05 | 0.02 | 0.05 |
| **ETA_Q** | 0.1 | 0.01 (0.01, 0.01) | 0.05 | 0.01 | 0.08 |
| **ETA_V2** | 0.08 | 0.02 (0.02, 0.02) | 0.01 | 0.02 | 0.04 |

**Supplementary Figure S1**. Concentration–time profiles for all 50 virtual subjects, with grey lines and symbols indicating subjects without outliers, and red lines and symbols indicating subjects with outliers.

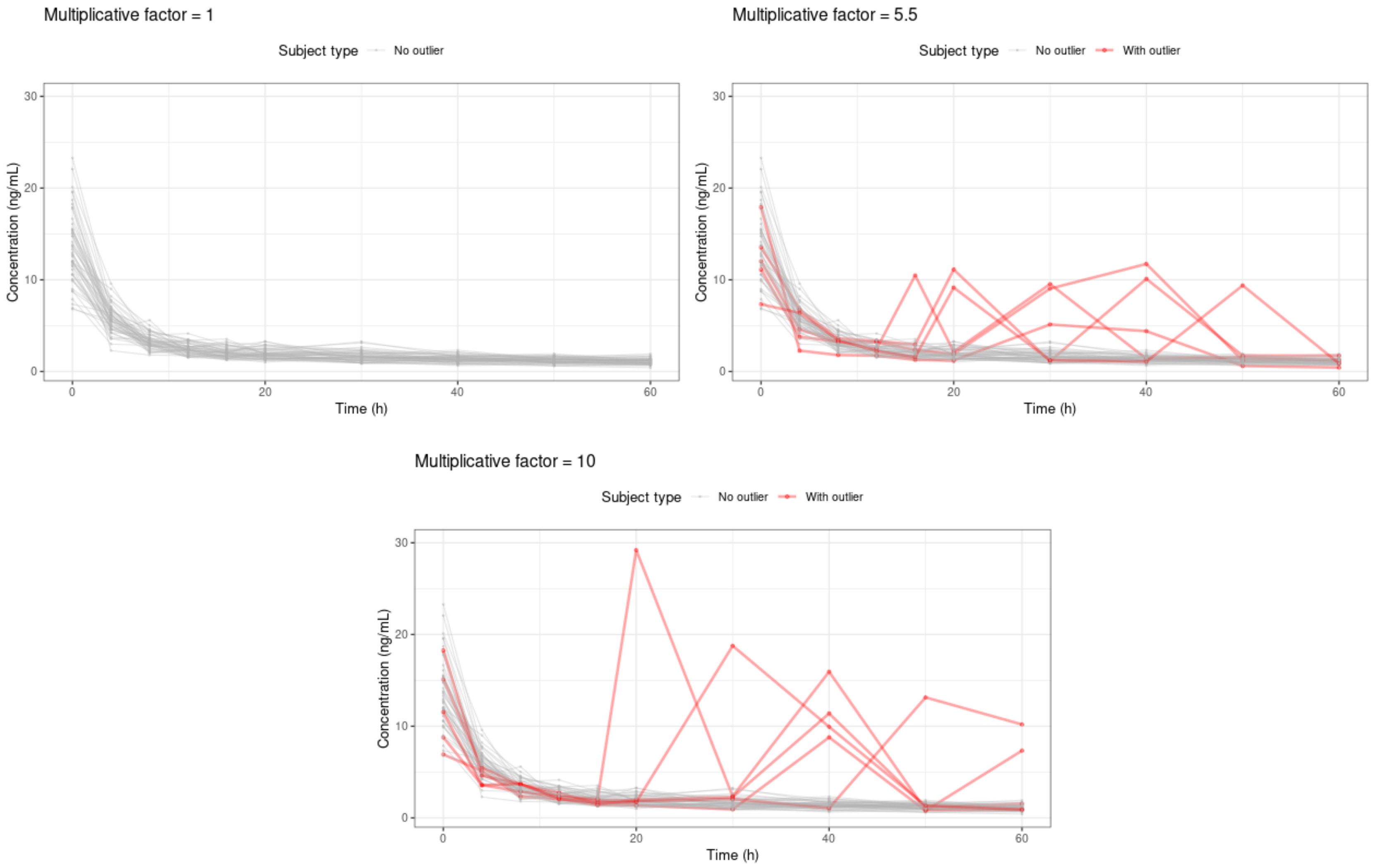

**Supplementary Figure S2**. Concentration–time profiles and corresponding posterior distributions for all 50 virtual subjects, shown by modeling method. Pink lines and shaded areas represent the posterior means and 90% credible intervals of the predicted concentrations, while black lines and points represent the observed data.

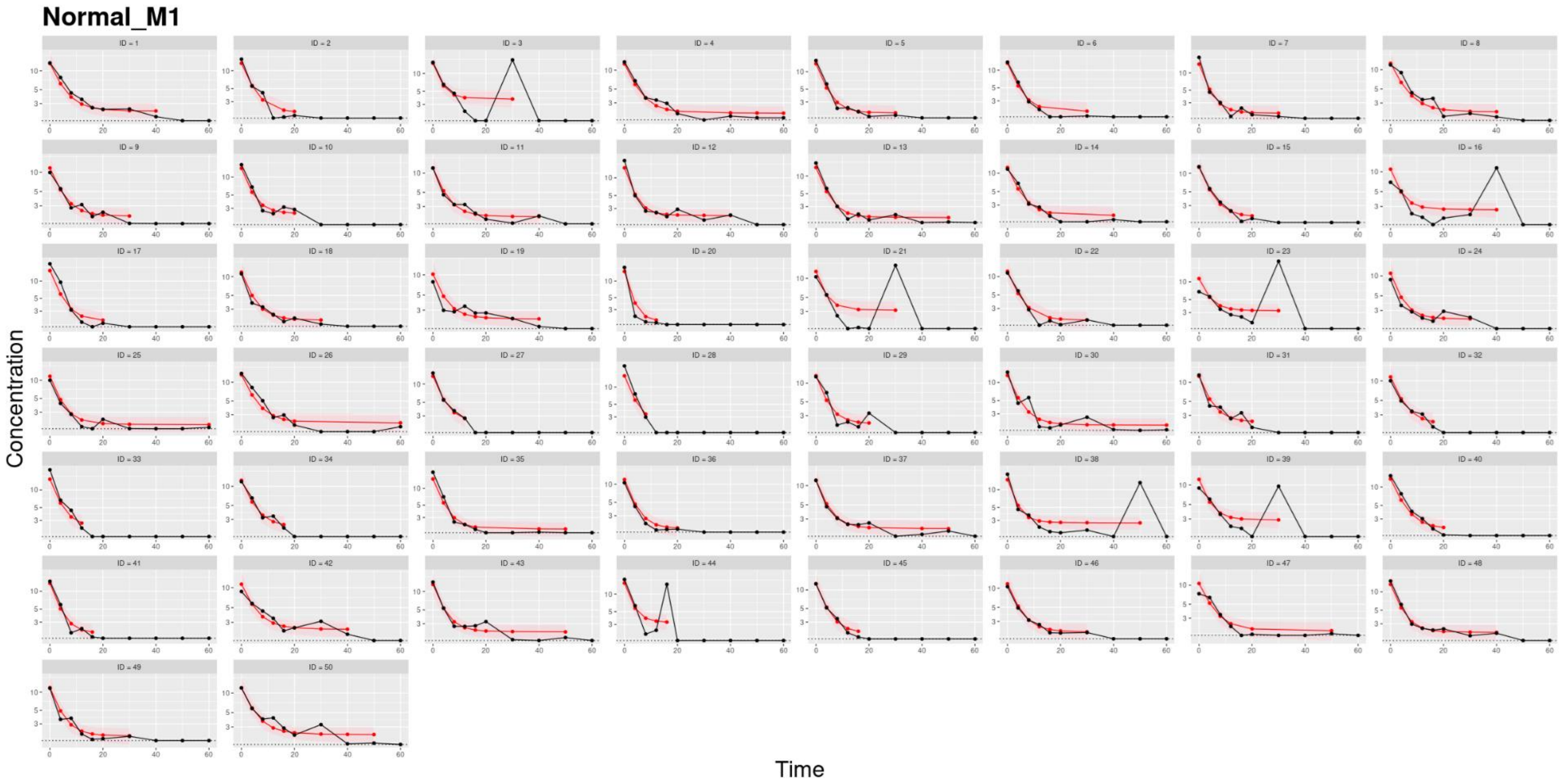

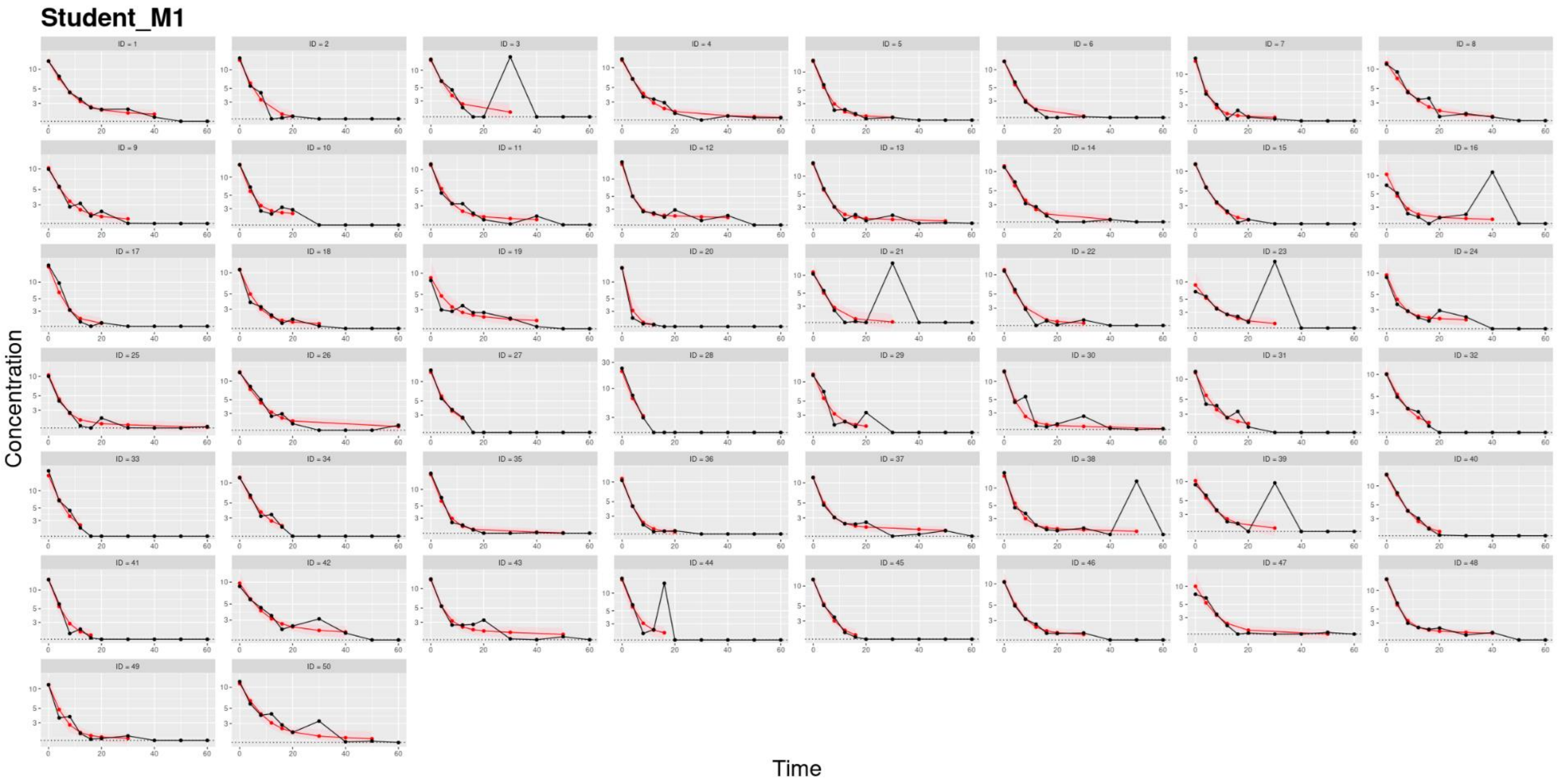
Student_M1
ID = 1
ID = 2
ID = 3
ID = 4
ID = 5
ID = 6
ID = 7
ID = 8
ID = 9
ID = 10
ID = 11
ID = 12
ID = 13
ID = 14
ID = 15
ID = 16
ID = 17
ID = 18
ID = 19
ID = 20
ID = 21
ID = 22
ID = 23
ID = 24
ID = 25
ID = 26
ID = 27
ID = 28
ID = 29
ID = 30
ID = 31
ID = 32
ID = 33
ID = 34
ID = 35
ID = 36
ID = 37
ID = 38
ID = 39
ID = 40
ID = 41
ID = 42
ID = 43
ID = 44
ID = 45
ID = 46
ID = 47
ID = 48
ID = 49
ID = 50
Concentration
Time

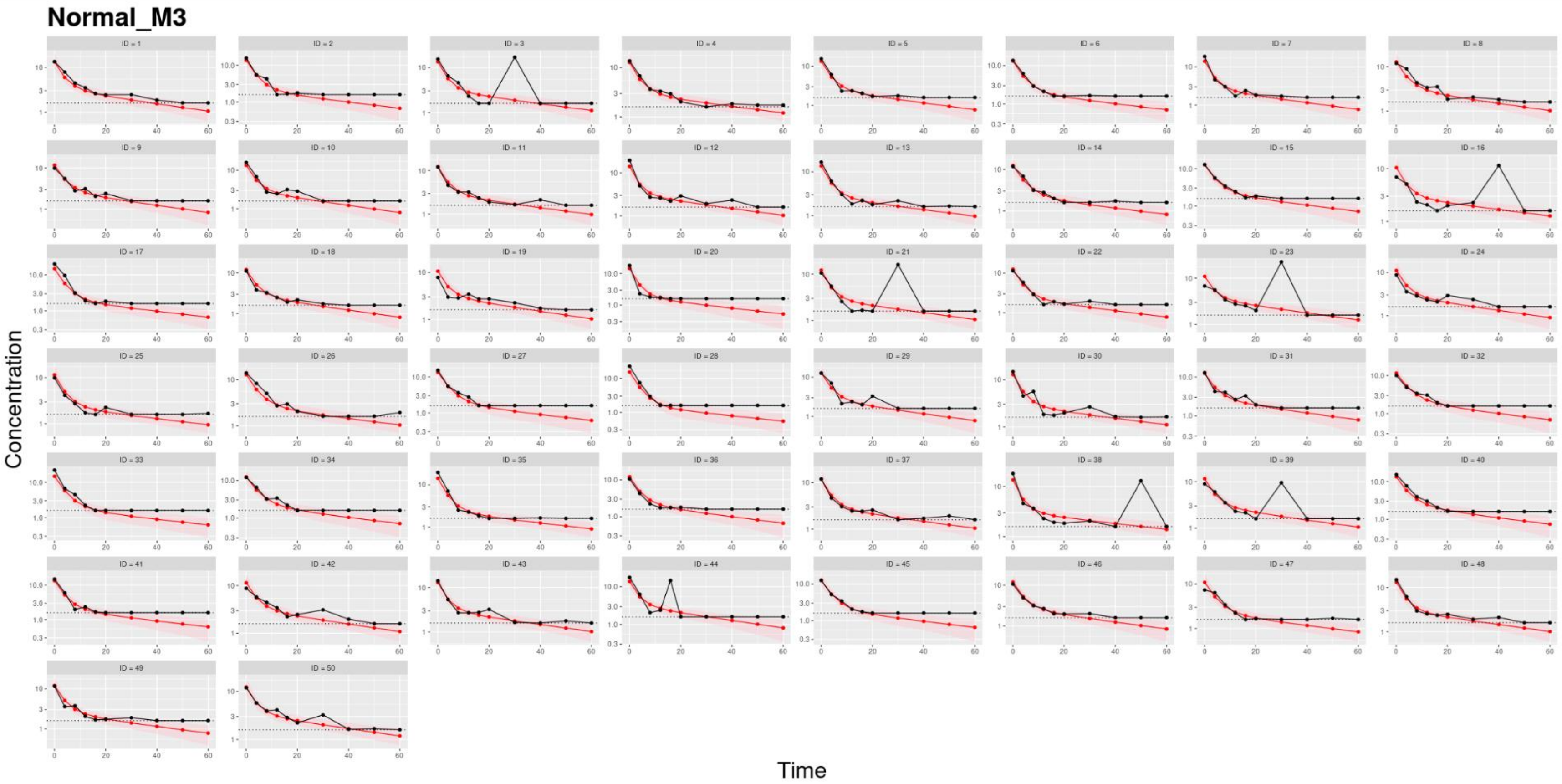
Normal_M3
ID = 1
ID = 2
ID = 3
ID = 4
ID = 5
ID = 6
ID = 7
ID = 8
ID = 9
ID = 10
ID = 11
ID = 12
ID = 13
ID = 14
ID = 15
ID = 16
ID = 17
ID = 18
ID = 19
ID = 20
ID = 21
ID = 22
ID = 23
ID = 24
ID = 25
ID = 26
ID = 27
ID = 28
ID = 29
ID = 30
ID = 31
ID = 32
ID = 33
ID = 34
ID = 35
ID = 36
ID = 37
ID = 38
ID = 39
ID = 40
ID = 41
ID = 42
ID = 43
ID = 44
ID = 45
ID = 46
ID = 47
ID = 48
ID = 49
ID = 50
Concentration
Time

### Student_M3

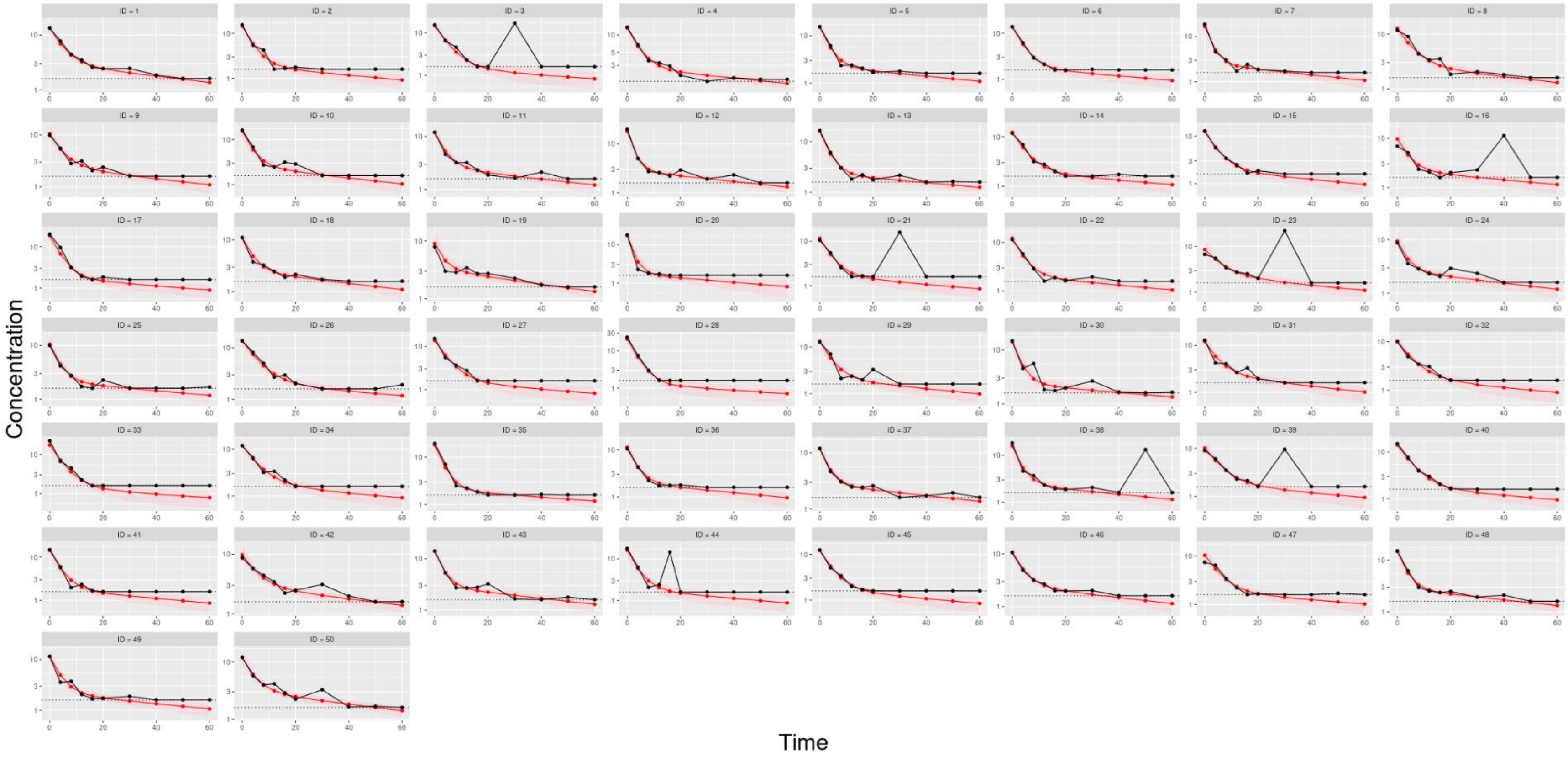

Concentration
Time
ID = 1
ID = 2
ID = 3
ID = 4
ID = 5
ID = 6
ID = 7
ID = 8
ID = 9
ID = 10
ID = 11
ID = 12
ID = 13
ID = 14
ID = 15
ID = 16
ID = 17
ID = 18
ID = 19
ID = 20
ID = 21
ID = 22
ID = 23
ID = 24
ID = 25
ID = 26
ID = 27
ID = 28
ID = 29
ID = 30
ID = 31
ID = 32
ID = 33
ID = 34
ID = 35
ID = 36
ID = 37
ID = 38
ID = 39
ID = 40
ID = 41
ID = 42
ID = 43
ID = 44
ID = 45
ID = 46
ID = 47
ID = 48
ID = 49
ID = 50

**Supplementary Figure S3**. Accuracy of individual PK parameter estimates across all subjects and four modeling methods used in the analysis underlying Table 1.

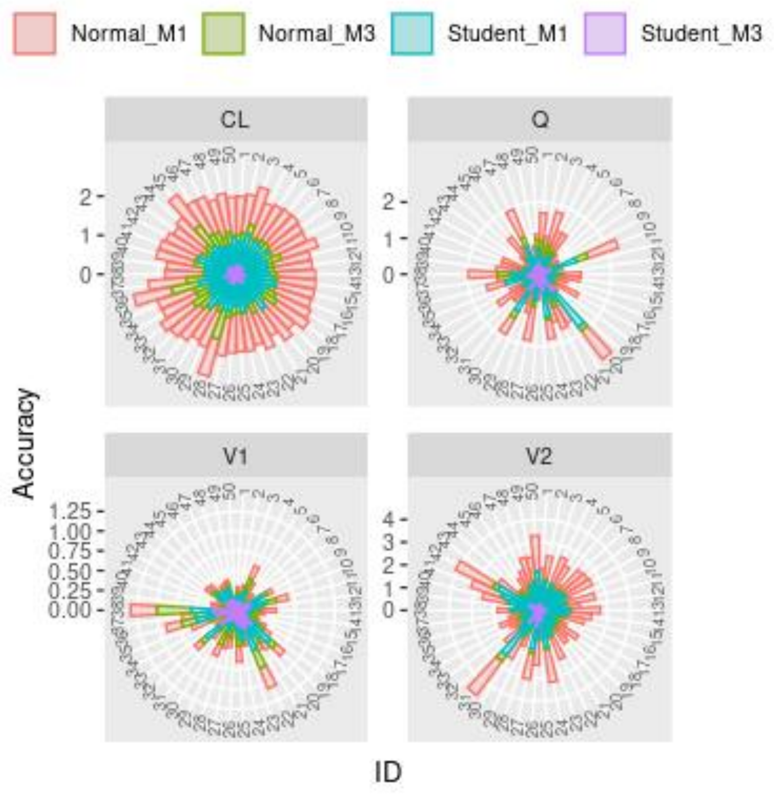

**Supplementary Figure S4.** Posterior distributions of population PK parameter estimates from the full Bayesian approach, based on a scenario in which 10% of subjects each had one outlier introduced at a concentration point within the α-phase. Dotted black lines indicate the true values.

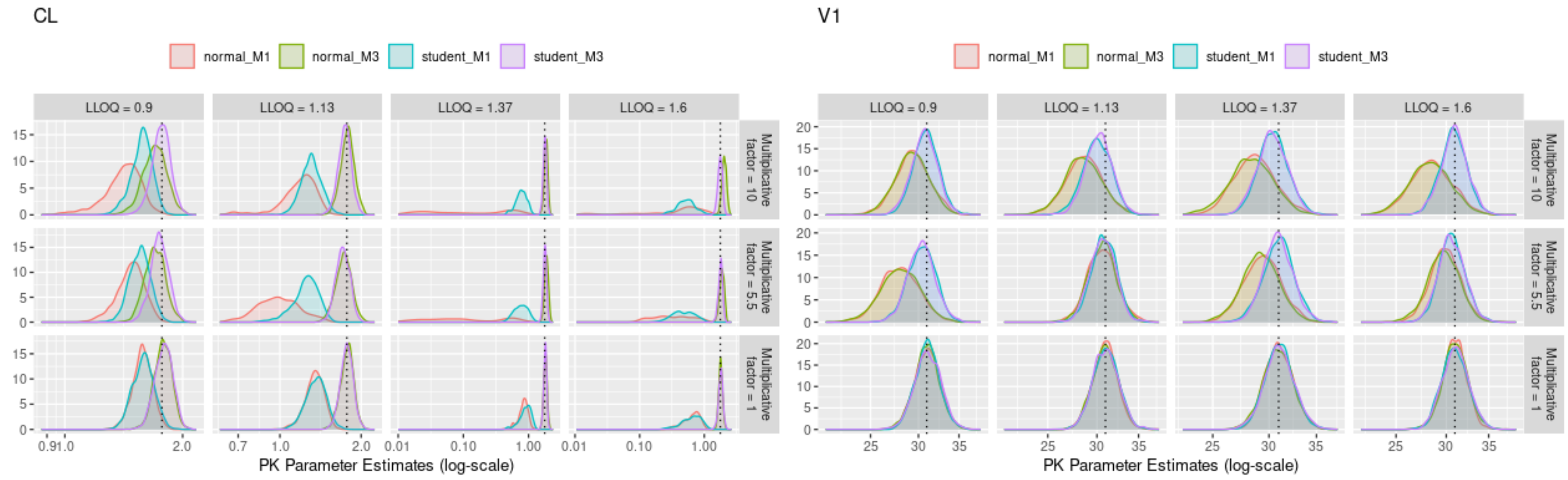

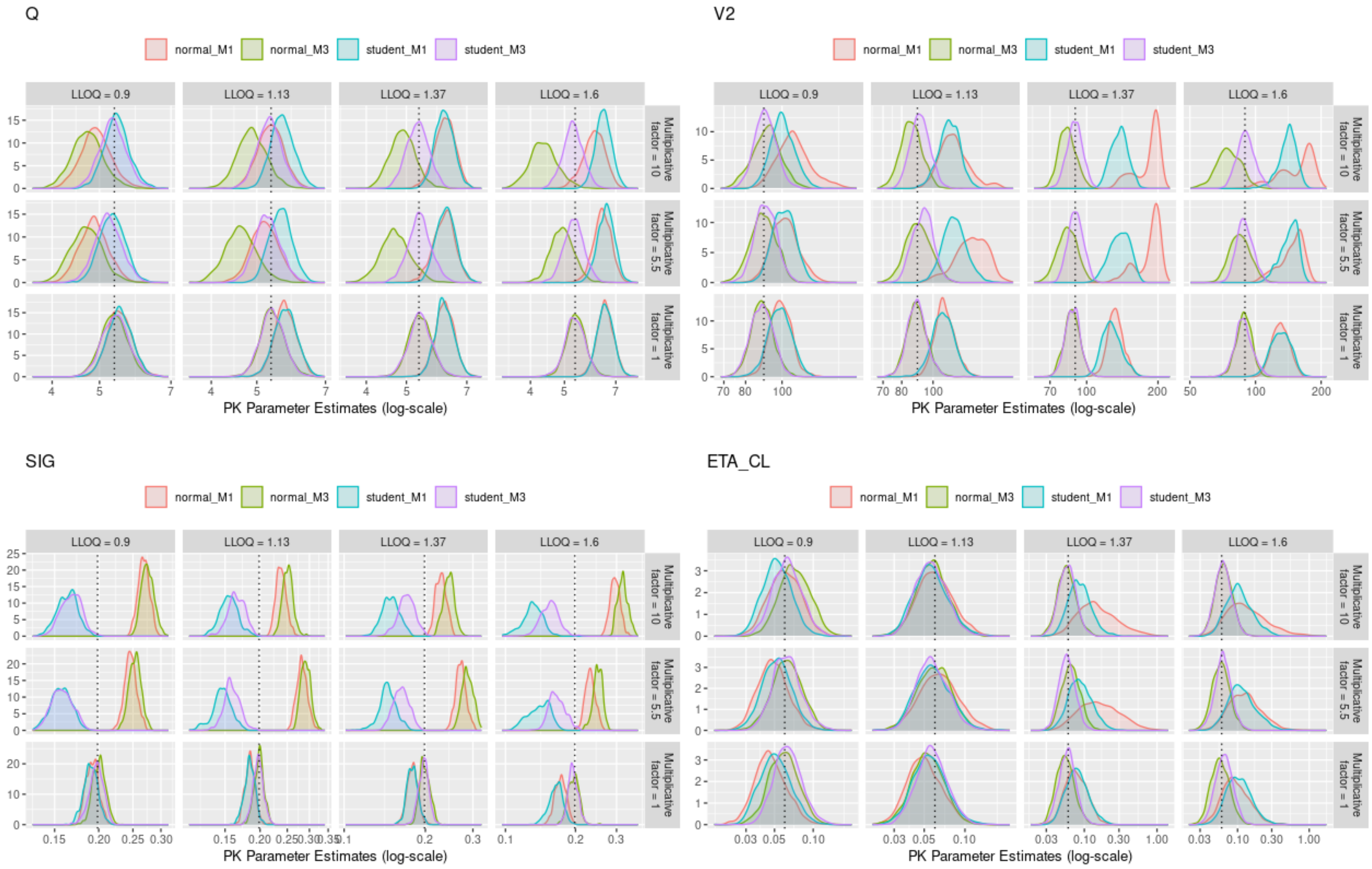
Q
V2
SIG
ETA_CL
normal_M1
normal_M3
student_M1
student_M3
LLOQ = 0.9
LLOQ = 1.13
LLOQ = 1.37
LLOQ = 1.6
Multiplicative factor = 10
Multiplicative factor = 5.5
Multiplicative factor = 1
PK Parameter Estimates (log-scale)

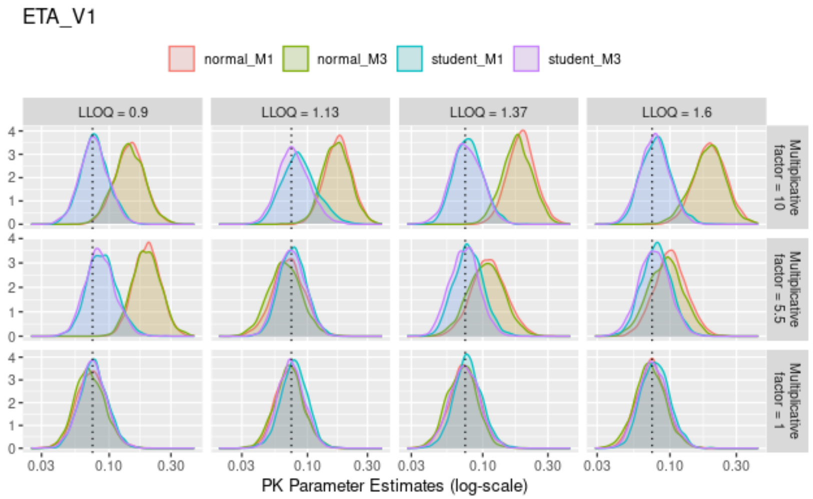
ETA_V1
normal_M1
normal_M3
student_M1
student_M3
LLOQ = 0.9
LLOQ = 1.13
LLOQ = 1.37
LLOQ = 1.6
Multiplicative factor = 10
Multiplicative factor = 5.5
Multiplicative factor = 1
PK Parameter Estimates (log-scale)

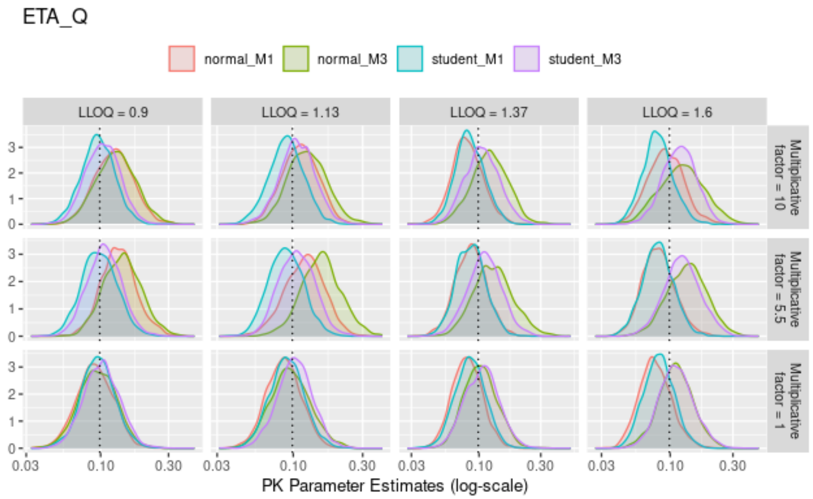
ETA_Q
normal_M1
normal_M3
student_M1
student_M3
LLOQ = 0.9
LLOQ = 1.13
LLOQ = 1.37
LLOQ = 1.6
Multiplicative factor = 10
Multiplicative factor = 5.5
Multiplicative factor = 1
PK Parameter Estimates (log-scale)

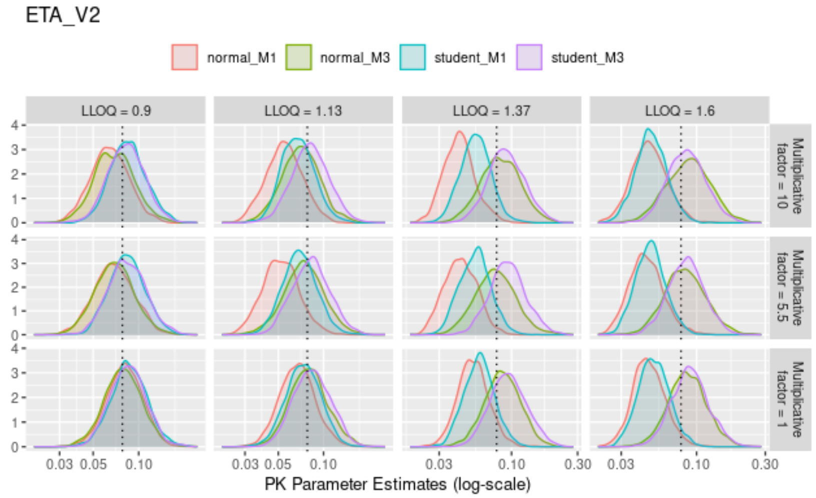
ETA_V2
normal_M1
normal_M3
student_M1
student_M3
LLOQ = 0.9
LLOQ = 1.13
LLOQ = 1.37
LLOQ = 1.6
Multiplicative factor = 10
Multiplicative factor = 5.5
Multiplicative factor = 1
PK Parameter Estimates (log-scale)

**Supplementary Figure S5.** Posterior distributions of population PK parameter estimates from the full Bayesian approach, based on a scenario in which 10% of subjects each had two outliers introduced: one concentration point from the α-phase and one from the β-phase. Dotted black lines indicate the true values.

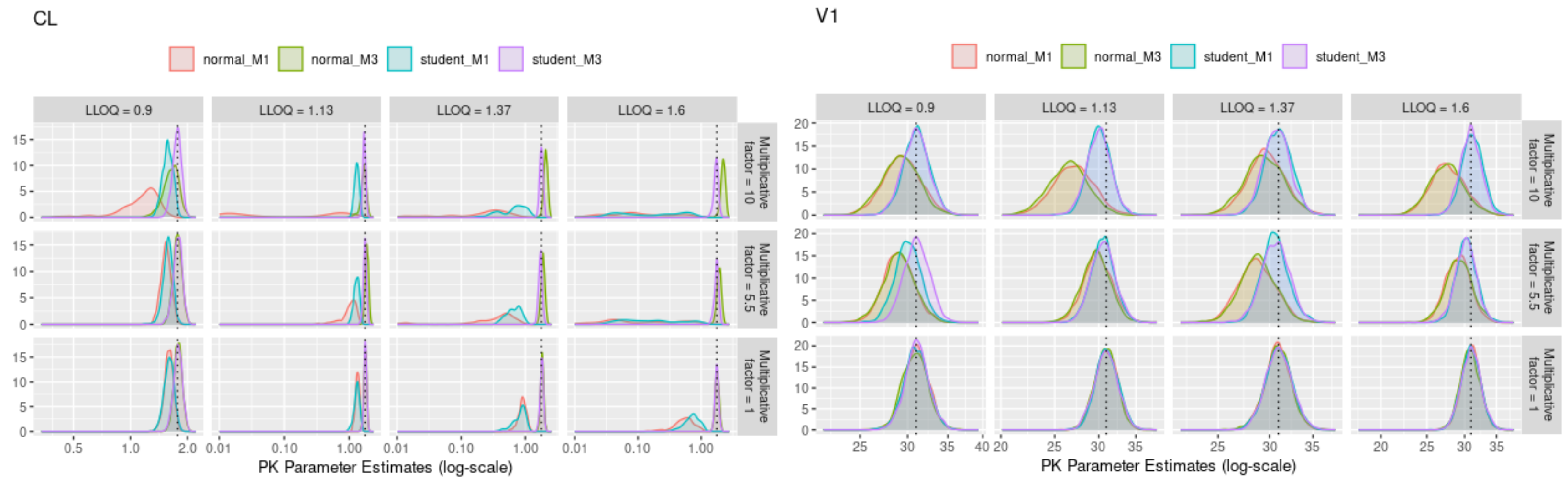

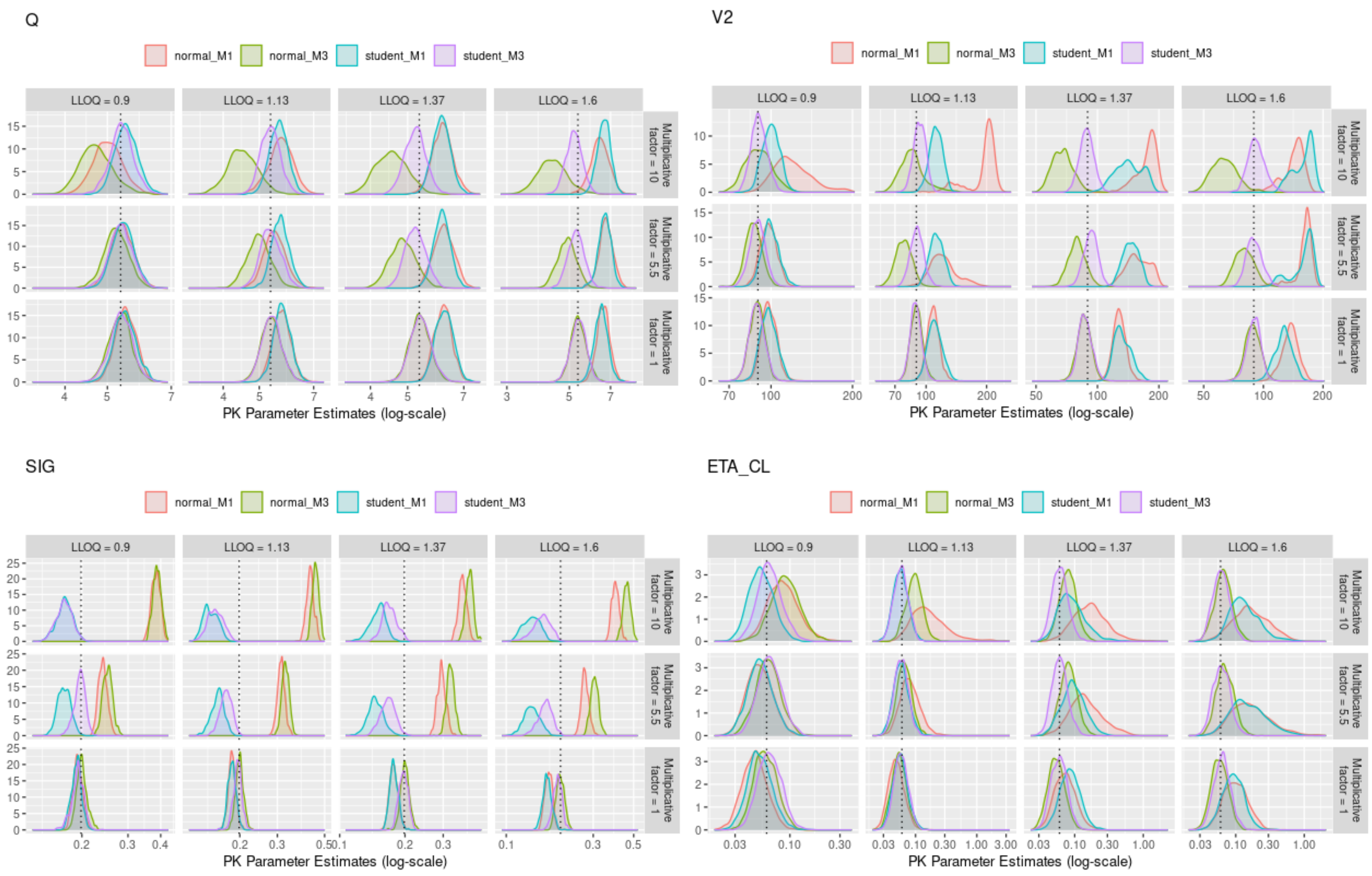

Q
V2
SIG
ETA_CL
normal_M1
normal_M3
student_M1
student_M3
LLOQ = 0.9
LLOQ = 1.13
LLOQ = 1.37
LLOQ = 1.6
Multiplicative factor = 10
Multiplicative factor = 5.5
Multiplicative factor = 1
PK Parameter Estimates (log-scale)

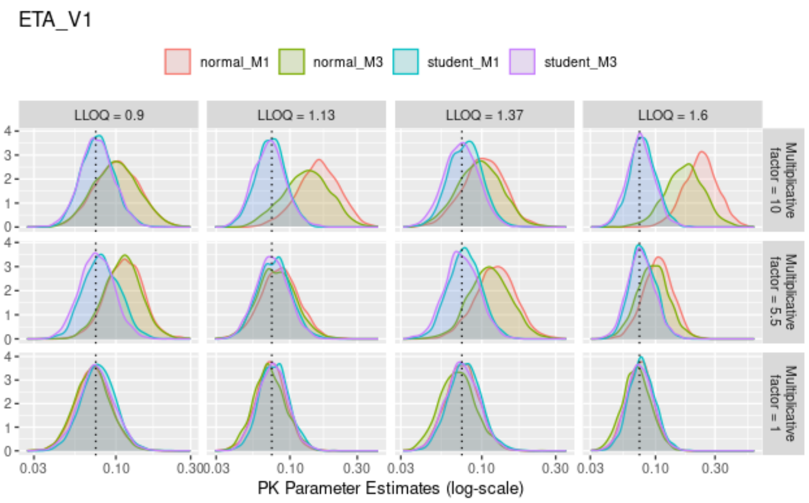
ETA_V1
normal_M1
normal_M3
student_M1
student_M3
LLOQ = 0.9
LLOQ = 1.13
LLOQ = 1.37
LLOQ = 1.6
Multiplicative factor = 10
Multiplicative factor = 5.5
Multiplicative factor = 1
0.03
0.10
0.30
PK Parameter Estimates (log-scale)

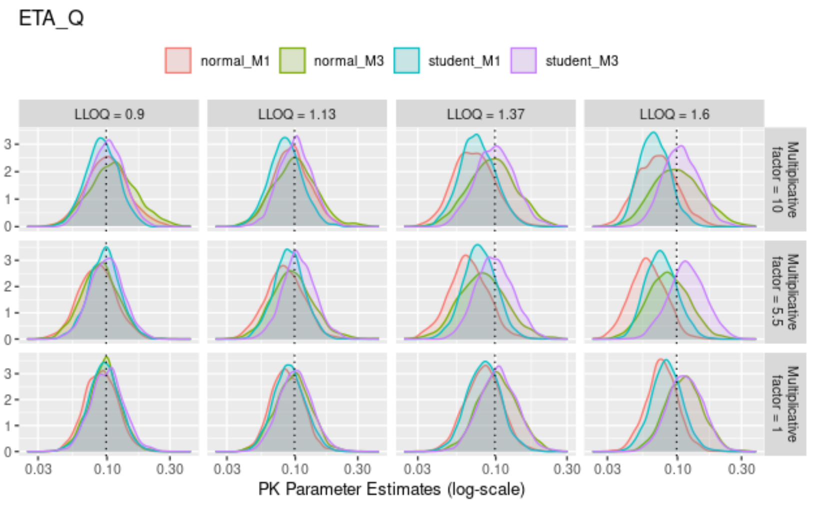
ETA_Q
normal_M1
normal_M3
student_M1
student_M3
LLOQ = 0.9
LLOQ = 1.13
LLOQ = 1.37
LLOQ = 1.6
Multiplicative factor = 10
Multiplicative factor = 5.5
Multiplicative factor = 1
0.03
0.10
0.30
PK Parameter Estimates (log-scale)

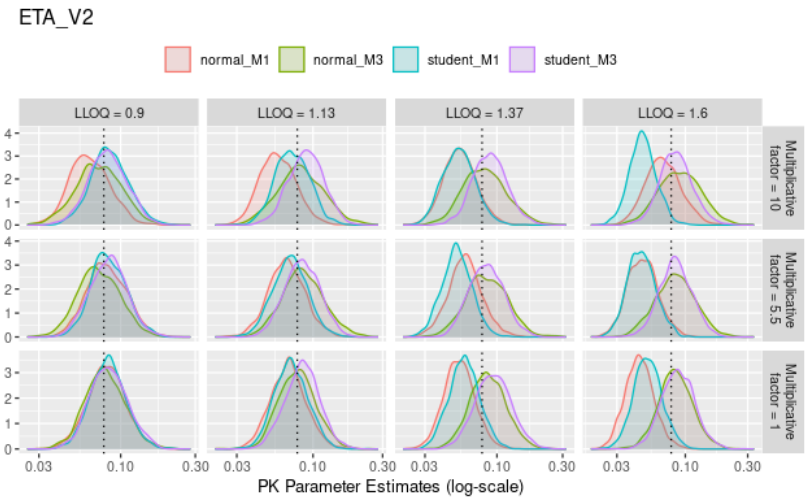
ETA_V2
normal_M1
normal_M3
student_M1
student_M3
LLOQ = 0.9
LLOQ = 1.13
LLOQ = 1.37
LLOQ = 1.6
Multiplicative factor = 10
Multiplicative factor = 5.5
Multiplicative factor = 1
0.03
0.10
0.30
PK Parameter Estimates (log-scale)

**Supplementary Figure S6.** Posterior distributions of population PK parameter estimates from the full Bayesian approach, based on a scenario in which 10% of subjects each had two outliers introduced from concentration points within the α-phase. Dotted black lines indicate the true values.

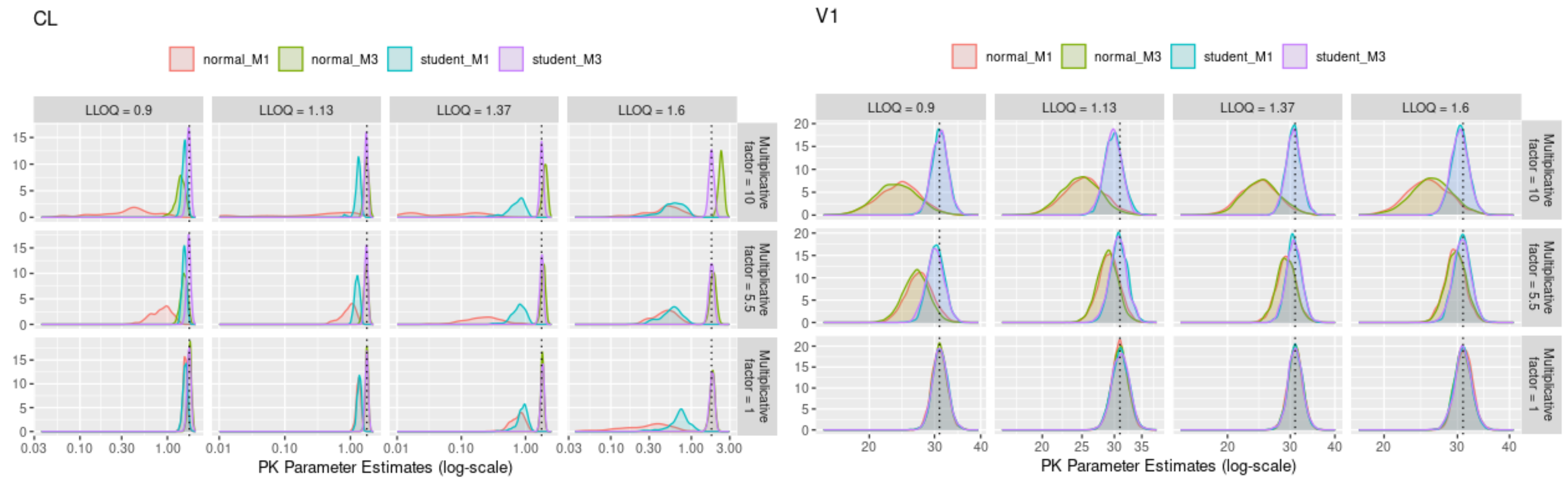

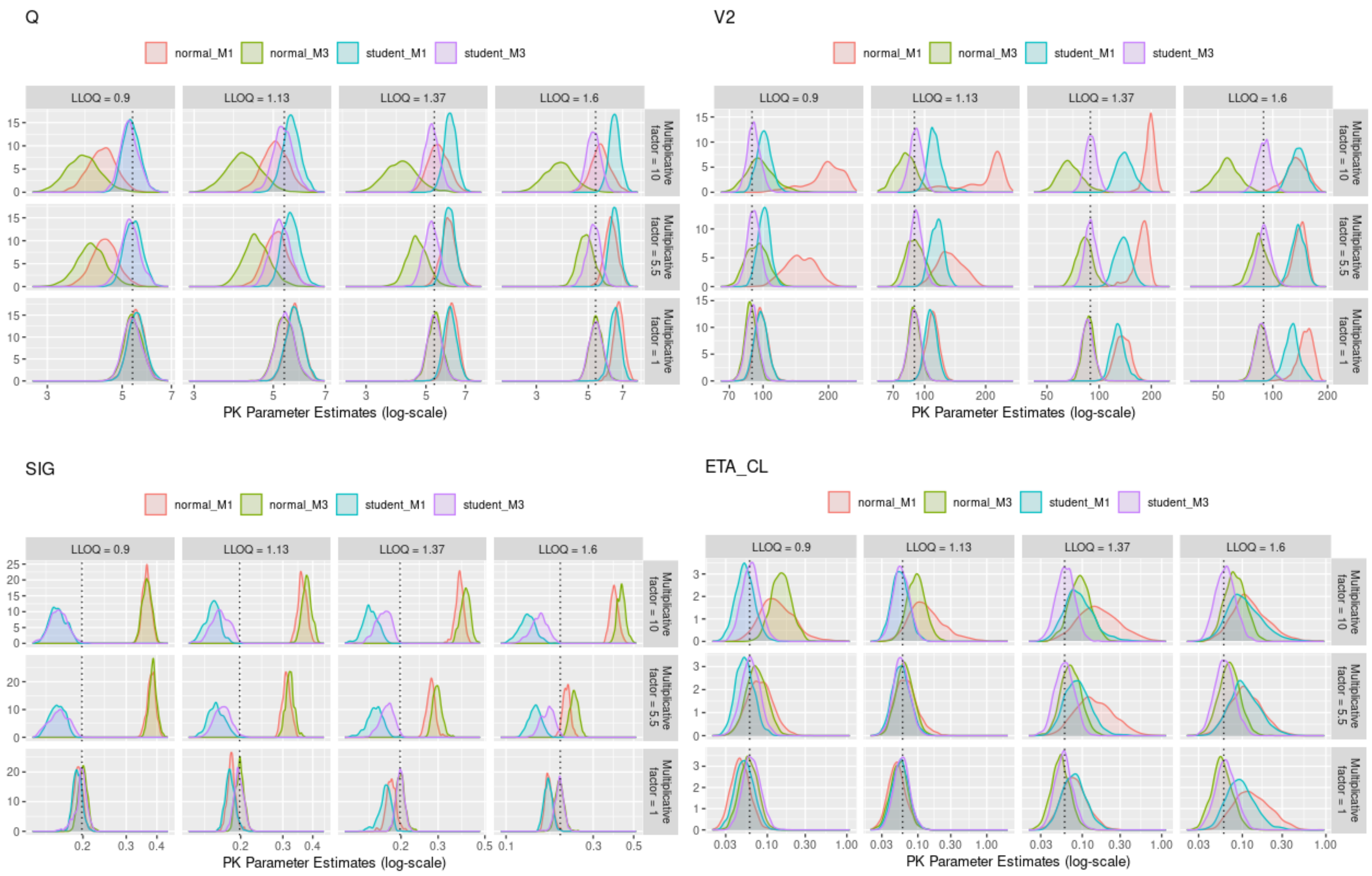

Q
V2
SIG
ETA_CL
normal_M1
normal_M3
student_M1
student_M3
LLOQ = 0.9
LLOQ = 1.13
LLOQ = 1.37
LLOQ = 1.6
Multiplicative factor = 10
Multiplicative factor = 5.5
Multiplicative factor = 1
PK Parameter Estimates (log-scale)

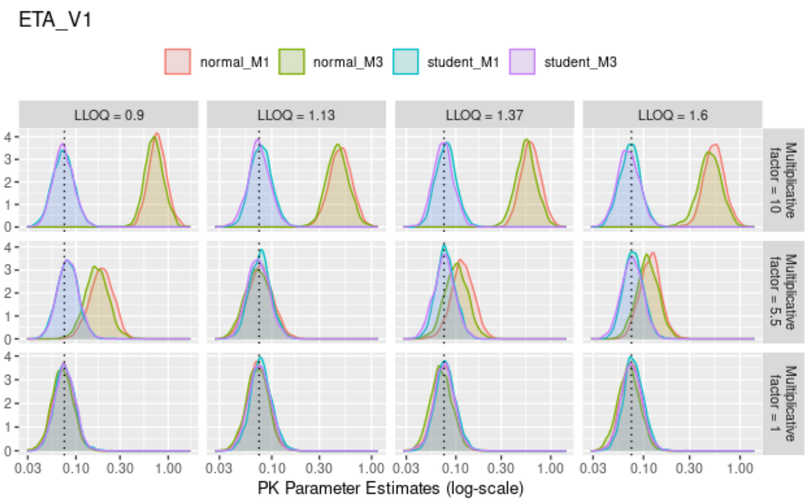
ETA_V1
normal_M1
normal_M3
student_M1
student_M3
LLOQ = 0.9
LLOQ = 1.13
LLOQ = 1.37
LLOQ = 1.6
Multiplicative factor = 10
Multiplicative factor = 5.5
Multiplicative factor = 1
PK Parameter Estimates (log-scale)

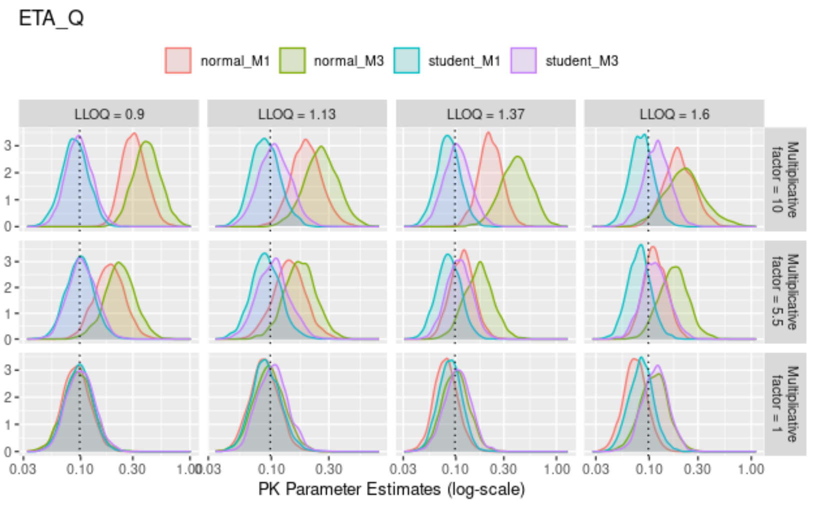
ETA_Q
normal_M1
normal_M3
student_M1
student_M3
LLOQ = 0.9
LLOQ = 1.13
LLOQ = 1.37
LLOQ = 1.6
Multiplicative factor = 10
Multiplicative factor = 5.5
Multiplicative factor = 1
PK Parameter Estimates (log-scale)

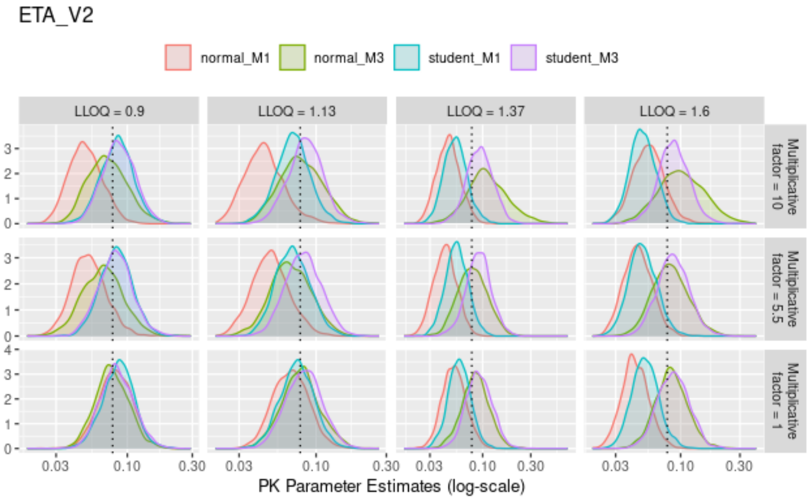
ETA_V2
normal_M1
normal_M3
student_M1
student_M3
LLOQ = 0.9
LLOQ = 1.13
LLOQ = 1.37
LLOQ = 1.6
Multiplicative factor = 10
Multiplicative factor = 5.5
Multiplicative factor = 1
PK Parameter Estimates (log-scale)

**Supplementary Figure S7.** Posterior distributions of population PK parameter estimates from the full Bayesian approach, based on a scenario in which 10% of subjects each had two outliers introduced from concentration points within the β-phase. Dotted black lines indicate the true values.

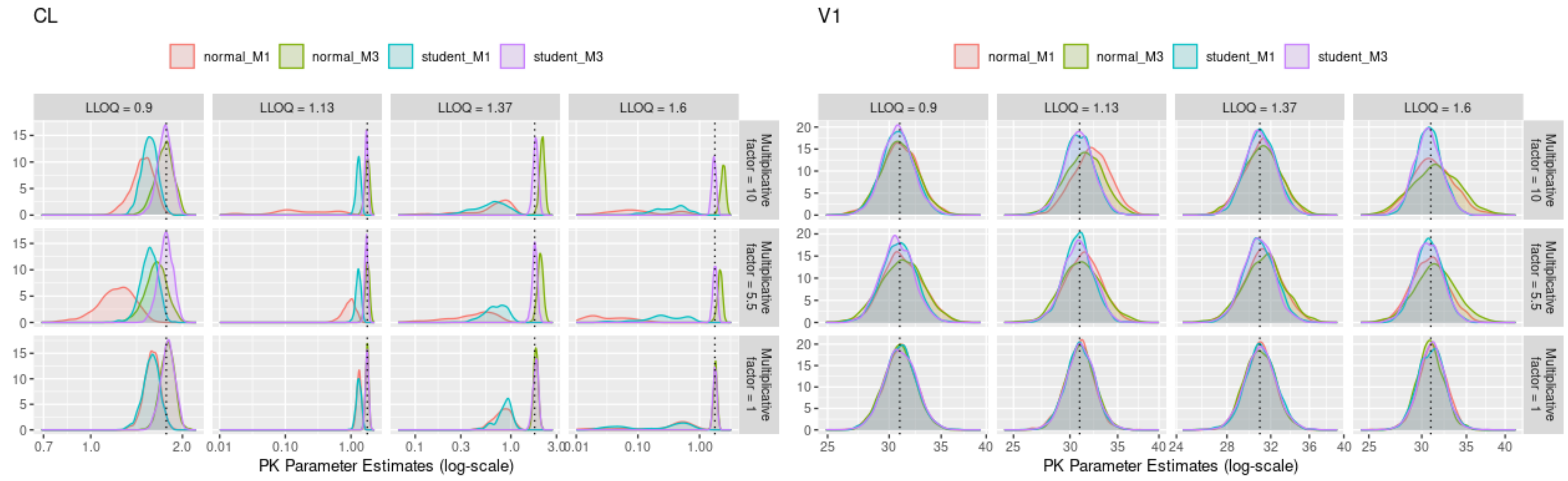

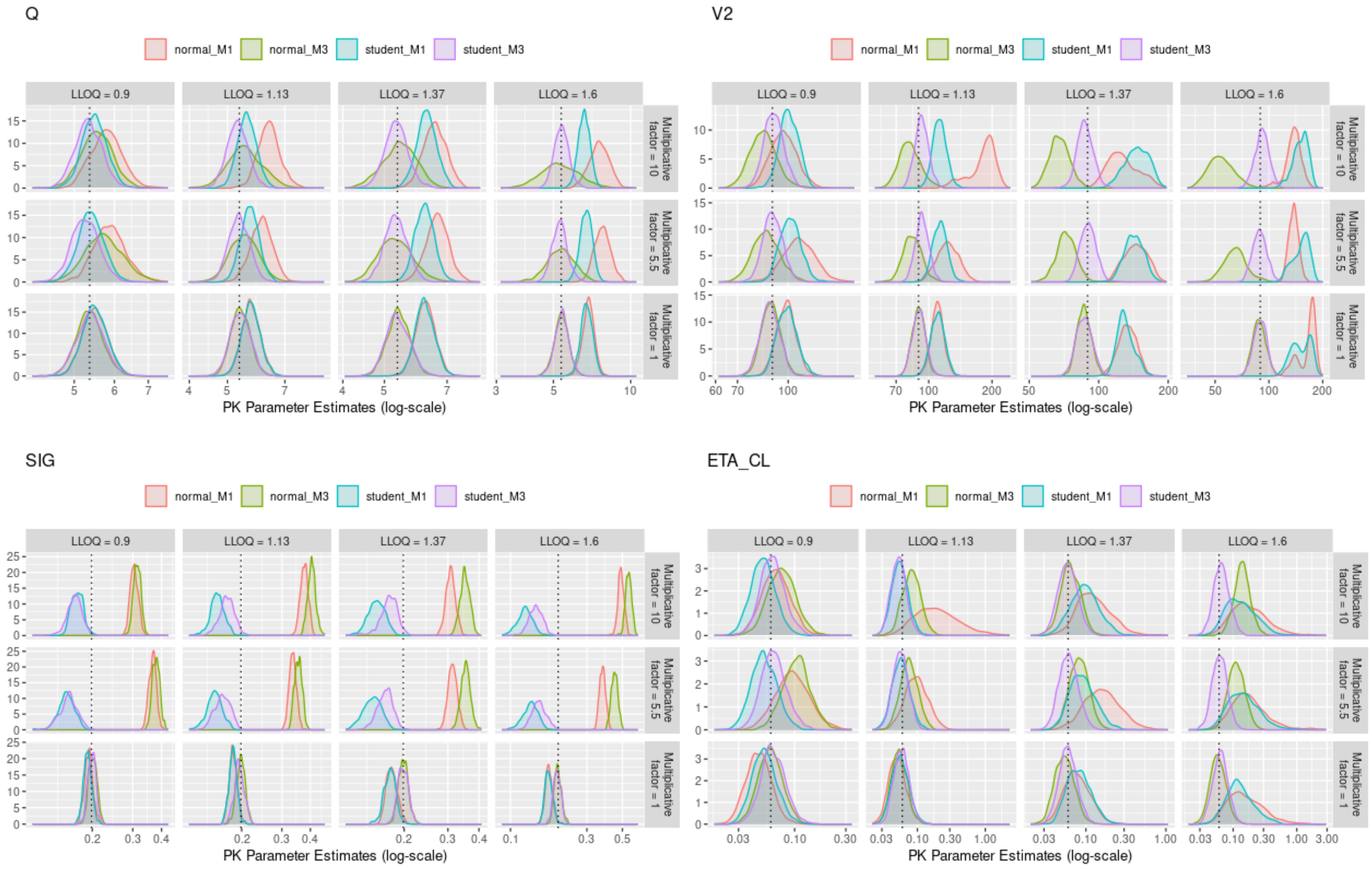
Q
V2
SIG
ETA_CL
normal_M1
normal_M3
student_M1
student_M3
LLOQ = 0.9
LLOQ = 1.13
LLOQ = 1.37
LLOQ = 1.6
Multiplicative factor = 10
Multiplicative factor = 5.5
Multiplicative factor = 1
PK Parameter Estimates (log-scale)

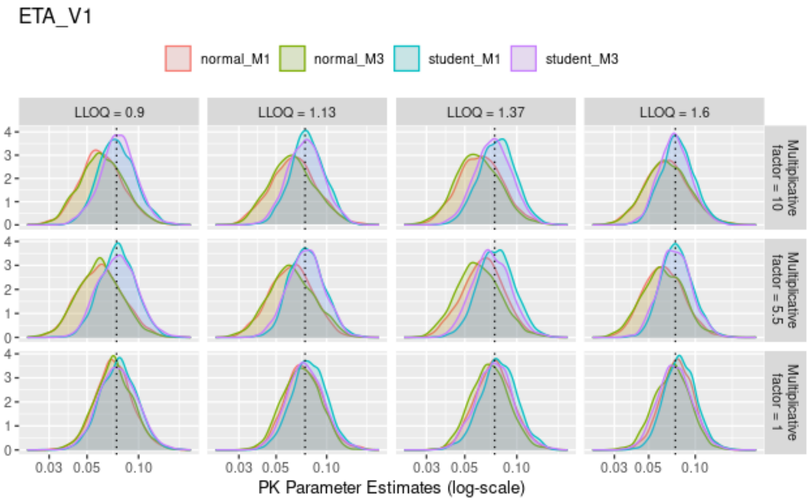
ETA_V1
normal_M1
normal_M3
student_M1
student_M3
LLOQ = 0.9
LLOQ = 1.13
LLOQ = 1.37
LLOQ = 1.6
Multiplicative factor = 10
Multiplicative factor = 5.5
Multiplicative factor = 1
0.03 0.05 0.10
PK Parameter Estimates (log-scale)

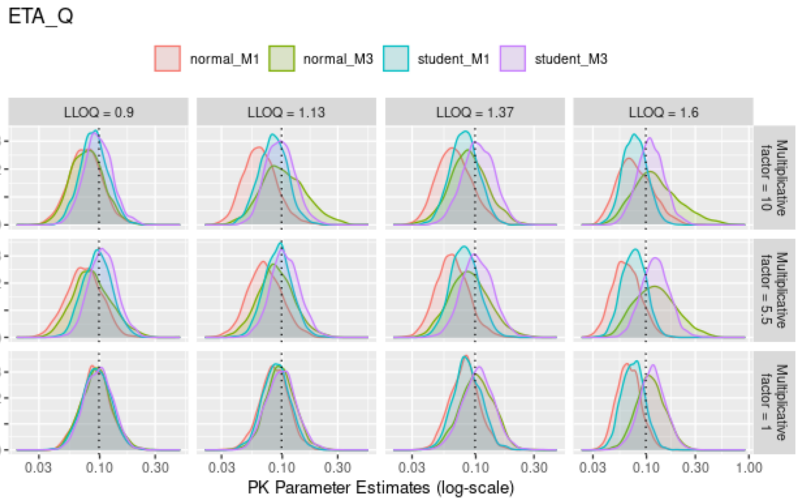
ETA_Q
normal_M1
normal_M3
student_M1
student_M3
LLOQ = 0.9
LLOQ = 1.13
LLOQ = 1.37
LLOQ = 1.6
Multiplicative factor = 10
Multiplicative factor = 5.5
Multiplicative factor = 1
0.03 0.10 0.30 1.00
PK Parameter Estimates (log-scale)

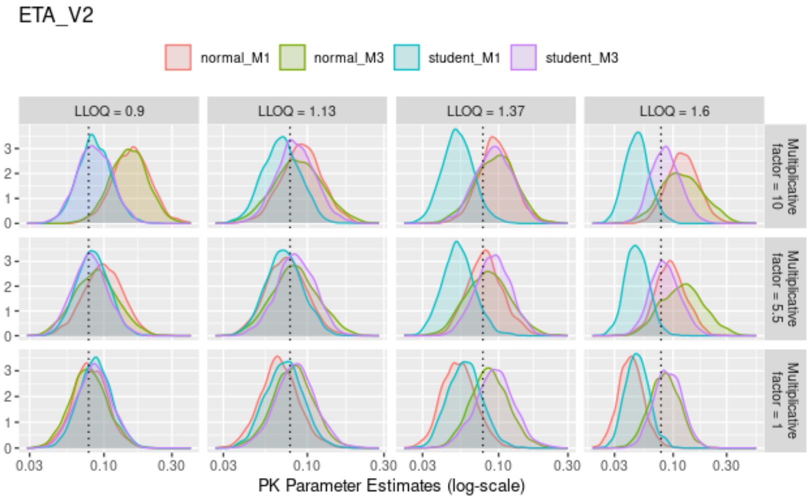
ETA_V2
normal_M1
normal_M3
student_M1
student_M3
LLOQ = 0.9
LLOQ = 1.13
LLOQ = 1.37
LLOQ = 1.6
Multiplicative factor = 10
Multiplicative factor = 5.5
Multiplicative factor = 1
0.03 0.10 0.30
PK Parameter Estimates (log-scale)
